# High dimensional co-expression networks enable discovery of transcriptomic drivers in complex biological systems

**DOI:** 10.1101/2022.09.22.509094

**Authors:** Samuel Morabito, Fairlie Reese, Negin Rahimzadeh, Emily Miyoshi, Vivek Swarup

## Abstract

Biological systems are immensely complex, organized into a multi-scale hierarchy of functional units based on tightly-regulated interactions between distinct molecules, cells, organs, and organisms. While experimental methods enable transcriptome-wide measurements across millions of cells, the most ubiquitous bioinformatic tools do not support systems-level analysis. Here we present hdWGCNA, a comprehensive framework for analyzing co-expression networks in high dimensional transcriptomics data such as single-cell and spatial RNA-seq. hdWGCNA provides built-in functions for network inference, gene module identification, functional gene enrichment analysis, statistical tests for network reproducibility, and data visualization. In addition to conventional single-cell RNA-seq, hdWGCNA is capable of performing isoform-level network analysis using long-read single-cell data. We showcase hdWGCNA using publicly available single-cell datasets from Autism spectrum disorder and Alzheimer’s disease brain samples, identifying disease-relevant co-expression network modules in specific cell populations. hdWGCNA is directly compatible with Seurat, a widely-used R package for single-cell and spatial transcriptomics analysis, and we demonstrate the scalability of hdWGCNA by analyzing a dataset containing nearly one million cells.

## Introduction

The development and widespread adoption of single-cell and spatial genomics approaches has lead to routine generation of high-dimensional datasets in a variety of biological systems. These technologies are frequently used to study developmental stages, evolutionary trajectories, disease states, drug perturbations, and other experimental conditions. Despite the inherent complexity and interconnectedness of biological systems, studies leveraging single-cell and spatial genomics typically analyze individual features (genes, isoforms, proteins, etc.) one by one, greatly oversimplifying the underlying biology. These datasets provide an opportunity for investigating and quantifying the relationships between these features to further contextualize their roles across biological conditions of interest.

Here we developed hdWGCNA, a framework for coexpression network analysis (1) in single-cell and spatial transcriptomics data. Co-expression networks are based on transformed pairwise correlations of input features, resulting in a quantitative measure of relatedness between genes (1, 2). Hierarchical clustering on the network structure allows us to uncover functional modules of genes whose expression profiles are tightly intertwined (3, 4), which typically correspond to specific biological processes and disease states. Considering that unique cell types and cell states have distinct gene expression programs, we designed hdWGCNA to facilitate multi-scale analysis of cellular and spatial hierarchies. hdWGCNA provides a rich suite of functions for data analysis and visualization, providing biological context for co-expression networks by leveraging a variety of biological knowledge databases. To maximize usability amongst the genomics community, the hdWGCNA R package extends the data structures and functionality of the widely-used Seurat package (5–7). We used hdWGCNA to analyze a single-cell RNA-seq (scRNA-seq) dataset consisting of 1M cells, showcasing the scalability of hdWGCNA in large datasets.

We applied hdWGCNA in a variety of high-dimensional transcriptomics datasets from different technologies and biological conditions. As a common use case, we performed iterative network analysis of the major cell types in the human prefrontal cortex (PFC), identifying shared and specific network modules in each cell type. We constructed co-expression networks in anterior and posterior mouse brain sections profiled with 10× Genomics Visium ST, and found distinct spatial patterns of these gene expression programs. Using long-read scRNA-seq (scRNA-seq) data from the mouse hippocampus (8), we uncovered splicing isoform co-expression networks in the radial glia lineage involved in cell fate specification. Network analysis of inhibitory neurons from published single-nucleus RNA-seq (snRNA-seq) in Autism spectrum disorder (ASD) donors (9) revealed modules disrupted in ASD containing key genetic risk genes such as *SCN2A, TSC1*, and *SHANK2*. We performed consensus co-expression network analysis of microglia from three Alzheimer’s disease (AD) snRNA-seq studies (10–12), yielding multiple gene modules corresponding to disease-associated microglia and polygenic risk of AD. Finally, we used hdWGCNA to project gene modules from the bulk RNA-seq AMP-AD cohort (13) into several published snRNA-seq datasets of AD brains, showing that our approach allows for interrogation of gene modules and networks that have been previously identified.

## Results

### Constructing co-expression networks from high-dimensional transcriptomics data

Here we describe a comprehensive framework for constructing and analyzing coexpression networks in high dimensional transcriptomic data (Fig. 1a). Given a gene expression dataset as input, coexpression network analysis typically consists of the following analysis steps: computing pairwise correlations of input features, weighting correlations with a soft power threshold (*β*), computing the topological overlap between features, and unsupervised clustering via the Dynamic Tree Cut algorithm (3) (Fig. S1, Methods). The sparsity and noise inherent in single-cell data can lead to spurious gene-gene correlations, thereby complicating co-expression network analysis. Additionally, the correlation structure of single-cell or spatial transcriptomic data varies greatly for different subsets (cell types, cell states, anatomical regions). A typical hdWGCNA workflow in scRNA-seq data accounts for these considerations by collapsing highly similar cells into “metacells” to reduce sparsity while retaining cellular heterogeneity, and by allowing for a modular design to perform separate network analyses in specified cell populations. Here, we demonstrate hdWGCNA in single-cell data through an iterative network analysis of six major cell types in a published dataset of human PFC samples from healthy donors (Fig. 1a) (11).

**Fig. 1.**
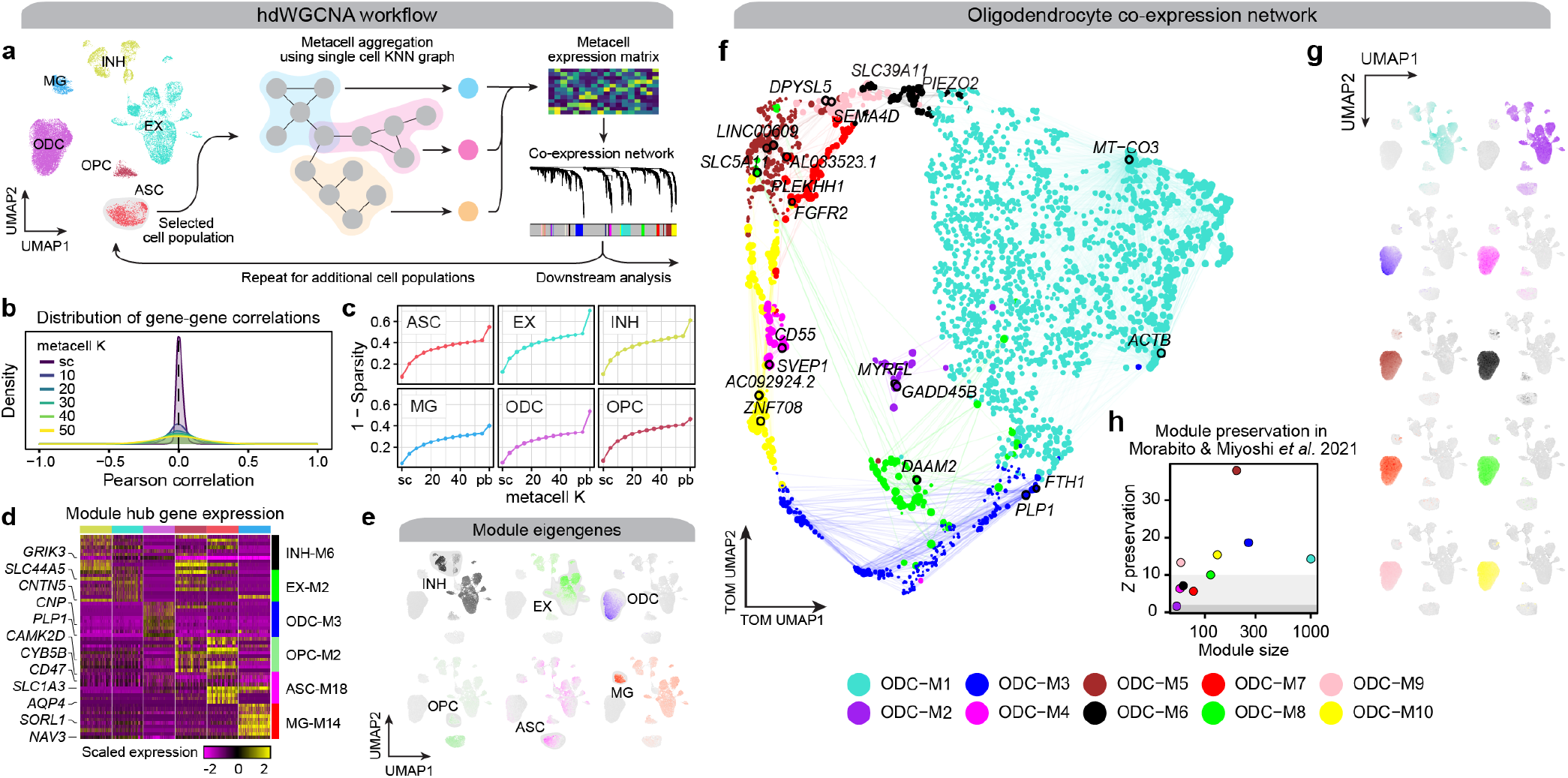
Overview of the hdWGCNA workflow and application in the human prefrontal cortex. **a.** Schematic overview of the standard hdWGCNA workflow on a scRNA-seq dataset. UMAP plot shows 36,671 cells from control donors in the Zhou *et al*. human PFC dataset. Cell type abbreviations are the following: ASC: astrocytes; EX: excitatory neurons; INH: inhibitory neurons; MG: microglia; ODC: oligodendrocytes; OPC: oligodendrocyte progenitor cells. **b.** Density plot showing the distribution of pairwise Pearson correlations between genes from the single-cell (sc) expression matrix, and metacell expression matrices with varying values of the *K*-nearest neighbors parameter *K*. **c.** Expression matrix density (1 - sparsity) for the single-cell (sc), pseudo-bulk (pb), and metacell matrices with varying values of *K* in each cell type. **d.** Heatmap of scaled gene expression for the top five hub genes by kME in INH-M6, EX-M2, ODC-M3, OPC-M2, ASC-M18, and MG-M14. **e.** snRNA-seq UMAP colored by module eigengene (ME) for selected modules as in **d**. **f.** UMAP plot of the ODC co-expression network. Each node represents a single gene, and edges represent co-expression links between genes and module hub genes. Point size is scaled by kME. Nodes are colored by co-expression module assignment. The top two hub genes per module are labeled. Network edges were downsampled for visual clarity. **g.** snRNA-seq UMAP as in **a** colored by MEs for the ten ODC co-expression modules as in **f**. **h.** Module preservation analysis of the ODC modules in the Morabito & Miyoshi et *al*. 2021 human PFC dataset. The module’s size versus the preservation statistic (*Z* preservation) is shown for each module. *Z* < 5: not preserved; 10 > *Z* ≥ 5: moderately preserved; *Z* ≥ 10: highly preserved.

Metacells are defined as small groups of transcriptomically similar cells representing distinctive cell states. There are several approaches to identify metacells from single-cell genomics data (14–17). Here, we leverage a bootstrapped aggregation (bagging) algorithm for constructing metacell transcriptomic profiles from single-cell datasets by applying K-nearest neighbors (KNN) to a dimensionality-reduced representation of the input dataset (Methods, Algorithm 1). This approach can be performed for each biological replicate to ensure that critical information about each sample (age, sex, disease status, etc.) is retained for downstream analysis. We computed gene-gene correlations in the normalized gene expression matrix from the single-cell dataset and metacell expression matrices while varying the number of cells to collapse into a single metacell (the KNN *K* parameter). The distribution of these gene-gene correlations displays a spike at zero for the single-cell expression matrix, with flattened distributions corresponding to more non-zero correlations in the metacell matrices, indicating that metacell expression profiles are less prone to noisy gene-gene correlations compared to the single-cell matrix (Fig. 1b) (Methods). We note that sparsity (defined in Equation 1) is greatly reduced in the metacell matrices for each cell type compared to the singlecell matrices, with over a tenfold reduction in some cases (Fig. 1c). We applied hdWGCNA to a dataset of CD34+ hematopoietic stem and progenitor stem cells (17) using two additional metacell approaches (15, 17), and found that all approaches were suitable for downstream network analysis (Supplementary Fig. S2, Supplementary Table 1).

While co-expression modules consist of many genes, it is often convenient to summarize the expression of the entire module into a single metric. WGCNA uses module eigengenes (MEs), the first principal component of the module’s gene expression matrix, to describe the expression patterns of co-expression modules. hdWGCNA computes MEs using specific accommodations for high-dimensional data, allowing for batch correction and regression of continuous covariates (Methods, Algorithm 2). Optionally, hdWGCNA can use alternative gene scoring methods such as or UCell (18) or Seurat’s `AddModuleScore` function, and we show that these scores are correlated with MEs (Fig. S3).

We constructed metacells and performed co-expression network analysis for each major cell type in the human PFC dataset (11) using the standard hdWGCNA workflow, yielding distinct network structures and sets of gene modules (Supplementary Tables 2-3). Through differential module eigengene (DME) analysis, we found shared and distinct modules across different cell types (Supplementary Table 4, Methods), and we highlight specific modules from each cell type (Figs. 1d, e). The expression of module hub genes, which are highly connected members of the coexpression network ranked by eigengene-based connectivity (kME), tend to display cell-type specific patterns, such as the myelination genes *CNP* and *PLP1* in oligodendrocyte (ODC) module ODC-M3 (Fig.1d). However, some co-expression modules may correspond to cellular processes common to multiple cell-types, in which case the hub genes may be widely expressed. We inspected the MEs of selected celltype specific modules, and found that the overall expression patterns were similar to that of their constituent hub genes (Fig. 1d,e)

We demonstrate some of the downstream functionalities of hdWGCNA using the ODC co-expression network (Fig. 1f-h). For network visualization, we used UMAP (19) to embed the co-expression network TOM into a two-dimensional manifold, using the topological overlap of each gene with the top hub genes from each module as input features (Methods, Fig. 1f). We found that eight of the ten ODC modules were specifically expressed in ODC cells based on their MEs (Fig. 1g, Wilcoxon rank sum test Bonferroni-adjusted *P*-value < 0.05). Finally, we performed module preservation analysis (20) to test the reproducibility of these modules in an independent dataset (12) and found that all of the ODC-specific modules were significantly preserved (*Z*-summary preservation ≥ 5). In sum, these network analyses in the human PFC dataset shows the core capabilities of the hdWGCNA workflow, (Fig. S1). Finally, we performed a similar iterative network analysis on a peripheral blood mononuclear cell (PBMC) scRNA-seq dataset of nearly 1M cells, highlighting the scalability of hdWGCNA in large datasets (Fig. S4, Supplementary Table 5).

### Spatial co-expression networks represent regional expression patterns in the mouse brain

Spatial transcriptomics (ST) enables the investigation of biological patterns that might otherwise be hidden in other -omics technologies such as single-cell or bulk RNA-seq (21, 22). Here we used hdWGCNA to identify spatial co-expression network modules in the murine brain using a publicly available Visium transcriptomics dataset from 10× Genomics (Fig. 2a). Sequencing-based ST approaches like Visium yield transcriptome-wide gene expression profiles localized to individual “spots”, where a single spot likely contains multiple cells. Data sparsity is also inherent to the current generation of these technologies, therefore we propose a *metaspot* aggregation approach prior to network analysis (Fig. S5). Evenly spaced spots throughout the input ST slide are used as principal spots, with at least one other spot in between two principal spots. The transcriptomes of the principal spots and their direct neighbors are aggregated into metaspot expression profiles, containing at most seven ST spots (Fig. S6a). Similar to metacells in scRNA-seq, the sparsity of the metaspot expression matrix was reduced compared to the original ST matrix S6b), and the distribution of gene-gene correlations in the metaspot expression matrix were less concentrated at zero S6c). hdWGCNA is capable of processing any number of ST samples in the same co-expression network analysis by constructing metaspots separately for each sample.

**Fig. 2.**
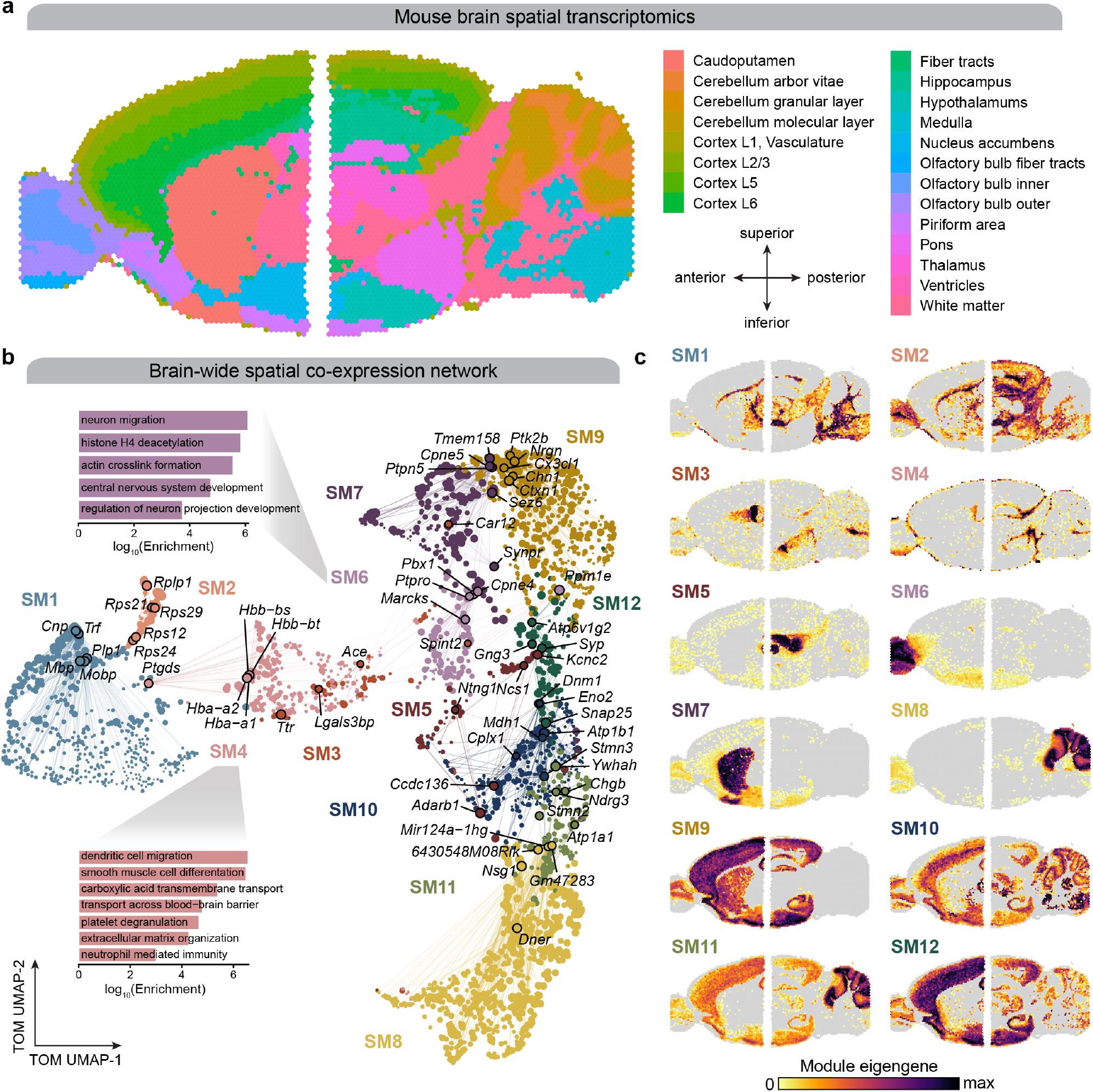
Spatial co-expression networks represent regional expression patterns in the mouse brain. **a.** Visium spatial transcriptomics (ST) in anterior (left) and posterior (right) mouse brain sections, colored by Louvain clusters annotated by anatomical regions. **b.** UMAP plot of the mouse brain ST co-expression network. Each node represents a single gene, and edges represent co-expression links between genes and module hub genes. Point size is scaled by kME. Nodes are colored by co-expression module assignment. The top 5 hub genes per module are labeled. Network edges were downsampled for visual clarity. **c.** ST samples colored by module eigengenes (MEs) for the 12 spatial co-expression modules. Grey color indicates a ME value less than zero.

We applied hdWGCNA in the mouse brain Visium dataset, identifying twelve spatial modules (SM1-12, Fig. S6, Supplementary Table 6), and we embedded the co-expression network in two dimensions using UMAP (Fig. 2b). DME analysis showed that spatial co-expression modules displayed distinct regional expression profiles based on their MEs (Fig. 2c, Supplementary Table 7), encompassing a wide array of cellular processes such as the myelination module SM1 in the white matter tracts, and synaptic transmission modules SM7, SM9, SM11, and SM12 (Fig. S6c, Supplementary Table 8). For example, DME analysis showed that expression of SM4 was localized to the ventricles and cortical layer 1 near the blood-brain barrier (Fig. 2c). Further, the hub genes of SM4 include hemoglobin subunits (*Hba-a1, Hba-a2, Hbb-bt*), and we show that SM4 was enriched for biological processes associated with brain vasculature (Figs. 2b, S6c). We compared these gene modules to cluster marker genes from a whole mouse brain snRNA-seq dataset (23) and found significant correspondences, such as the striatum module SM7 and medium spiny neurons (Fisher’s exact test FDR < 0.05, Fig. S6d). Additionally, we performed network analysis on a subset of this dataset containing cortical layers 2-6 (Fig. S7), identifying additional fine-grained spatial co-expression modules localized to specific cortical layers (Supplementary Tables 9, 10).

### Isoform-level co-expression networks reveal cell fate decisions in the radial glia developmental lineage

Different isoforms of the same gene are often involved in distinct biological processes (26). Conventional single-cell transcriptomics assays capture information at the gene level, thereby missing much of the biological diversity and regulatory mechanisms that occurs at the isoform level (27). Emerging long-read sequencing approaches enable us to profile cellular transcriptomes at isoform resolution (8, 28–30), thus providing new opportunities to model the relationships between isoforms using co-expression network analysis.

We used hdWGCNA to perform isoform co-expression network analysis in radial glia lineage cells from the mouse hippocampus at postnatal day 7 (P7) profiled with ScISOrSeq (8) (Fig. 3a, Methods). Radial glia, which share transcriptomic similarities with mature astrocytes, are progenitor cells that give rise to numerous distinct cell fates including neuronal cells, astrocytes, oligodendrocytes, and ependymal cells (31, 32). To model this developmental process, we applied Monocle3 (33) pseudotime to 2,190 radial glia lineage cells (Fig. 3b). We identified three trajectories corresponding to distinct cell fates, termed the ependymal (EPD) trajectory, astrocyte (ASC) trajectory, and the neural intermediate progenitor cell (NPC) trajectory.

**Fig. 3.**
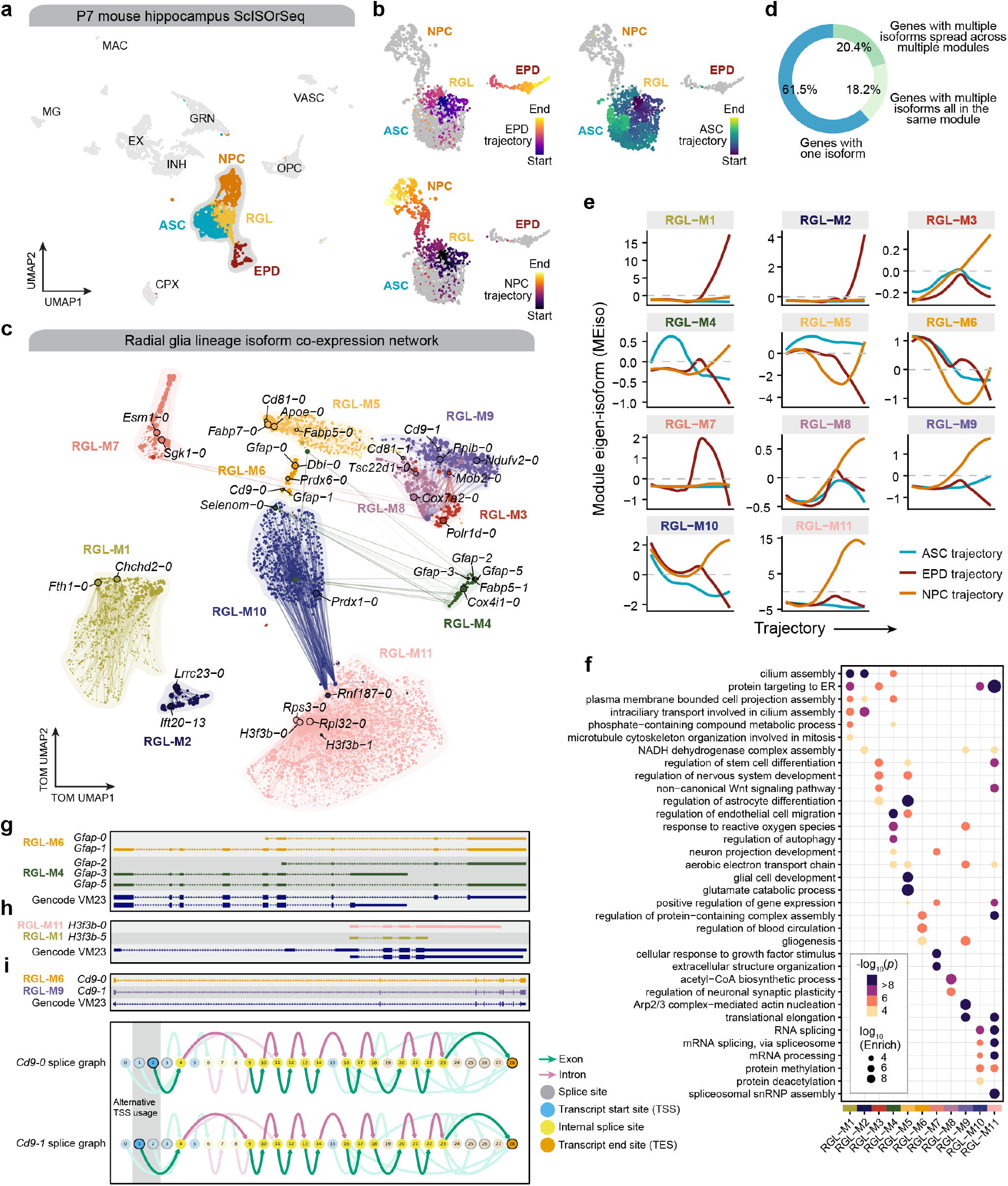
Co-expression network analysis of the radial glia lineage in the mouse hippocampus. **a.** UMAP plot of cells from the mouse hippocampus ScISOrSeq dataset (8). Major cell types are labeled and the cells used for co-expression network analysis are colored. This dataset contains expression information for 96,093 isoforms and 31,053 genes in 6,832 cells. **b.** UMAP plot of the radial glia lineage, colored by Monocle 3 (24) pseudotime assignment. Top left: ependymal (EPD) trajectory; top right: astrocyte (ASC) trajectory; bottom left: neuronal intermediate progenitor cell (NPC) trajectory. **c.** UMAP plot of the radial glia lineage isoform co-expression network. Each node represents a single isoform, and edges represent co-expression links between isoforms and module hub isoforms. Point size is scaled by kMEiso. Nodes are colored by co-expression module assignment. Network edges were downsampled for visual clarity. **d.** Donut chart showing the percentage of genes with one isoform, with multiple isoforms that are all assigned to the same module, and with multiple isoforms that are spread across more than one module. **e.** Module eigenisoforms (MEiso) as a function of pseudotime for each co-expression module. For each module, a separate loess regression line is shown for each developmental trajectory. **f.** Dot plot showing selected GO term enrichment results for each co-expression module. **g.** Gene models for selected isoforms of *Gfap*, colored by co-expression module assignment. **h.** Gene models for selected isoforms of *H3f3b*, colored by co-expression module assignment. **i.** Top: gene models for selected isoforms of *Cd9*, colored by co-expression module assignment. Bottom: Swan (25) graphical representation of *Cd9* alternative splicing isoforms. Splice sites and transcript start / end sites are represented as nodes; introns and exons are represented as connections between nodes. These two isoforms are distinguished by alternative TSS usage. Gene models from the GENCODE VM23 comprehensive transcript set are shown below transcripts in panels **g**,**h**, and **i**.

Isoform co-expression network analysis revealed eleven modules in the radial glia lineage (Fig. 3c, Supplementary Table 11). Of the genes retained for network analysis, 61.5% had a single isoform, 18.2% had multiple isoforms that were all assigned to the same module, and 20.4% had multiple isoforms spread across several modules (Fig. 3d). Thus, these network modules capture information about the roles of different isoforms of the same gene in distinct biological processes. We inspected module eigenisoform (MEiso) patterns throughout the developmental lineage, thereby uncovering isoform modules critical for cell fate decisions (Fig. 3e, Supplementary Tables 12, 13). Increased expression of modules RGL-M1 and RGL-M2, which were enriched cilium assembly genes (Fig. 3f), was associated with the transition from a radial glia to an ependymal cell state. A steady expression level of module RGL-M5 (glial development, astrocyte differentiation) was found in the transition from radial glia to astrocytes, while a decreased expression of RGL-M5 lead to alternative fates. Four modules (RGL-M3, RGL-M8, RGL-M9, and RGL-M11) displayed an increase in expression in the neuronal trajectory, containing genes associated with cellular processes such as non-canonical *Wnt* signaling, neuronal synaptic plasticity, and RNA splicing (Fig. 3f).

We inspected the isoforms of three selected genes which had hub isoforms in different co-expression modules: *Gfap, H3f3b*, and *Cd9* (Figs. 3g-i). *Gfap* encodes a key intermediate filament protein in astrocytes that is involved in astrocytic reactivity during central nervous system injuries or neurodegeneration (34), and we found that modules RGL-M4 and RGL-M6 contained hub isoforms of *Gfap* featuring alternative splicing, alternative transcription start site (TSS) usage, and alternative transcription end site (TES) usage (Fig. 3g). Different isoforms of the histone H3.3 subunit gene *H3f3b* were hubs for modules RGL-M1 and RGL-M11, which were associated with ependymal and neuronal cell fates respectively, suggesting that alternative TES usage in *H3f3b* plays a role in regulating epigenetic factors in murine hippocampal development (Fig. 3f). *Cd9* encodes a transmembrane protein and is a known glioblastoma biomarker (35), and we found subtle differences in the TSS between hub isoforms in modules RGL-M6 and RGL-M9 that we show as a splicing summary graph (25) (Fig. 3i), supporting functional changes mediated by small isoform differences.

### Co-expression network analysis of inhibitory neurons in Autism spectrum disorder

Co-expression networks can be interrogated to further understand the molecular phenotypes of complex polygenic diseases in primary human tissue samples. We applied hdWGCNA to inhibitory neruons (INH) from a published snRNA-seq dataset of the PFC in Autism spectrum disorder (ASD) patients and age-matched controls (9) (Figs. 4a, S9, Supplementary Table 14). The INH network contained 14 modules, and we show hub genes that have a known association with ASD in the SFARI database on the co-expression UMAP (Fig. 4b). The MEs showed that some modules were primarily confined to a single INH cluster (INH-M3, INH-M1) while others were spread across multiple neuronal groups (Fig. 4c). Furthermore, DME analysis revealed significant differences between MEs in ASD and control samples for all modules except INH-M4 in at least one INH subpopulation (Fig. 4d, Supplementary Table 15, Wilcoxon rank sum test Bonferroni-adjusted *P*-value < 0.05). Three co-expression modules (INH-M11, INH-M13, and INH-M3) were significantly enriched in ASD-associated genes from the SFARI database (Fig. 4e), but we note that all of these modules contained several ASD-associated SFARI genes.

**Fig. 4.**
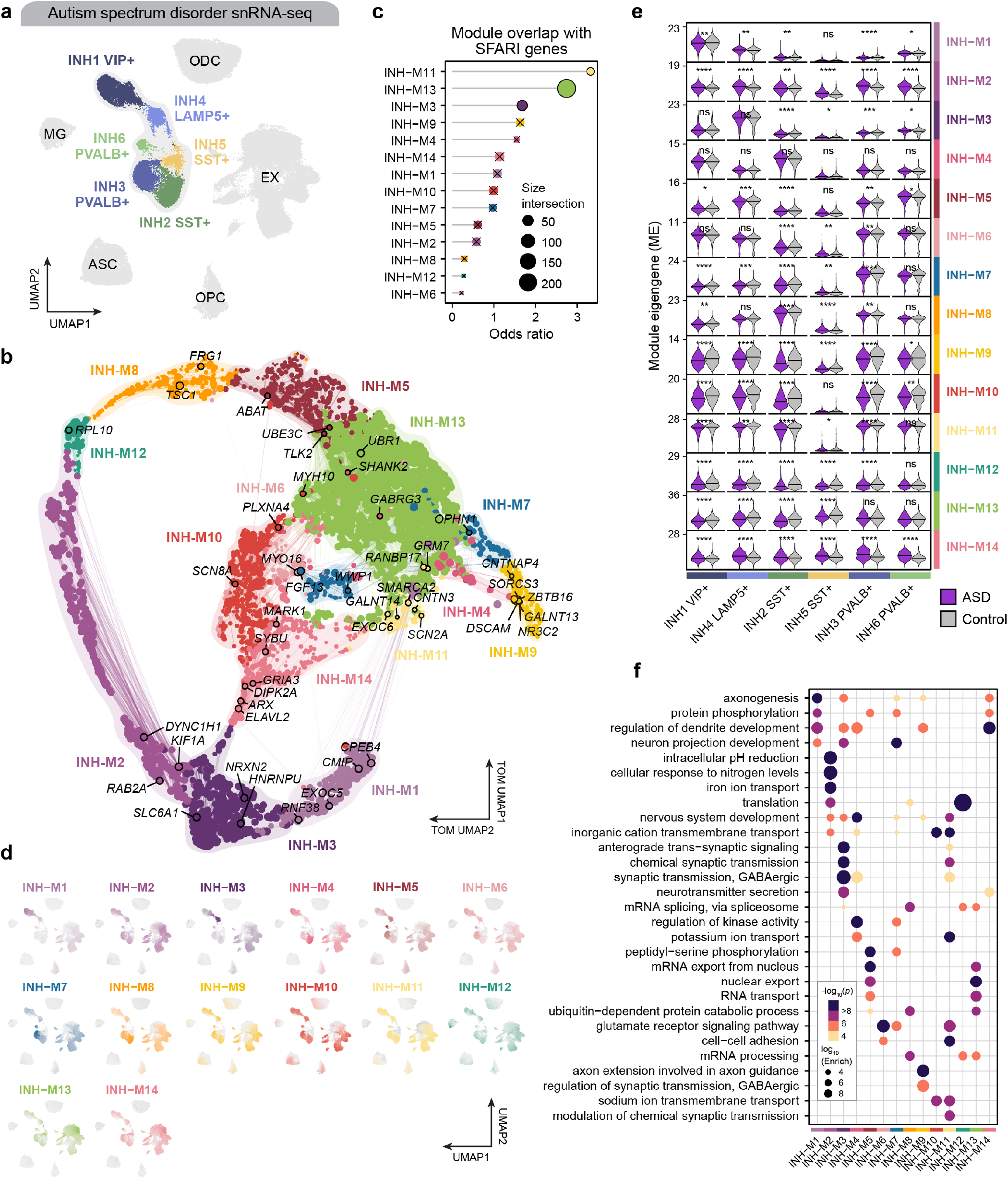
Co-expression network analysis of inhibitory neurons in Autism spectrum disorder. **a.** UMAP plot of 121,451 nuclei from the cortex of ASD donors and controls profiled with snRNA-seq. Inhibitory neuron subtypes are highlighted. **b.** Gene co-expression network derived from inhibitory neurons, represented as a two-dimensional UMAP embedding of the TOM. Nodes represent genes, colored by module assignment. Module hub genes with prior evidence of ASD association from SFARI are labeled. Edges represent co-expression relationships between genes and module hub genes. Network edges were downsampled for visual clarity. **c.** Gene overlap analysis comparing ASD-associated genes from SFARI and INH co-expression modules, using Fisher’s exact test. × indicates that the overlap was not significant (FDR > 0.05). **d.** snRNA-seq UMAP plots as in panel **A** colored by module eigengenes (MEs) for INH co-expression modules. **e.** Violin plots showing MEs in each INH cluster. Two-sided Wilcoxon test was used to compare ASD versus control samples. Not significant (ns): *P* > 0.05; *: *P* ≤ 0.05; **: *P* ≤ 0.01; ***: *P* ≤ 0.001; ****: *P* ≤ 0.0001. **f.** Selected GO enrichment results for each co-expression module.

INH-M11 was enriched for genes associated with synaptic transmission, ion transport, glutamate receptor signaling, and nervous system development (Fig. 4f, Supplementary Table 16), and this module was down-regulated in ASD for five of the six INH subtypes (Fig. 4d). Similarly, INH-M13 was associated with RNA processing (Fig. 4f) and was down-regulated in ASD in all INH subtypes except *PVALB+* neurons (Fig. 4d). One of the ING-M13 hub genes is *CHD2*, whose de novo variants have been identified in individuals with ASD (36, 37). CHD2 is part of the CHD family of chromatin modifying proteins and can alter gene expression by modification of chromatin structure. Similarly, rare loss-of-function mutations have been reported in *SCN2A* gene, a hub gene of INH-M11 module (38). We also find enrichment of several ASD-associated genes like *TSC1* (INH-M8), *SMARCA4* (INH-M8), *SHANK2* (INH-M4), *CPEB4* (INH-M1), highlighting that these modules are functional and provide new insights into the role of inhibitory neurons in ASD. Finally, we tested for the preservation of these modules in a snRNA-seq dataset from of the PFC from major depressive disorder (MDD) donors, and found substantial evidence of preservation across all modules except INH-M1 (Figs.S9c-e).

### Consensus network analysis of microglia in Alzheimer’s disease

Microglia, the resident immune cells of the brain, are implicated in the pathology and genetic risk of several central nervous system (CNS) diseases, including AD (39–42). Transcriptomic and epigenomic studies in human tissue and AD mouse models have identified multiple cell states of microglia, representing a spectrum between homeostatic and disease-associated microglia (DAMs) (12, 43, 44). Our previous study defined a set of transcription factors, genes, and *cis*-regulatory elements involved in the shift between homeostatic and DAM cell states in human AD, identifying shared and distinct signatures compared to the DAM signature from 5xFAD mice (12). Here we sought to expand on previous work by providing a systems level analysis of gene expression throughout the spectrum of microglia cell states.

We modeled the cell-state continuum between homeostatic and DAM-like microglia by employing a pseudotime analysis of microglia from three human AD snRNA-seq datasets (10–12) (Figs.5a, b, S10). Next, we performed consensus co-expression network analysis using microglia integrated from three human AD snRNA-seq datasets (10–12), identifying four consensus modules (Fig. 5c, Supplementary Table 17). Consensus network analysis is an approach that performs network analysis separately for each dataset, followed by a procedure to retain structures common across the individual networks, and thus it is well-suited for analyzing microglia co-expression from these different sources (Methods).

**Fig. 5.**
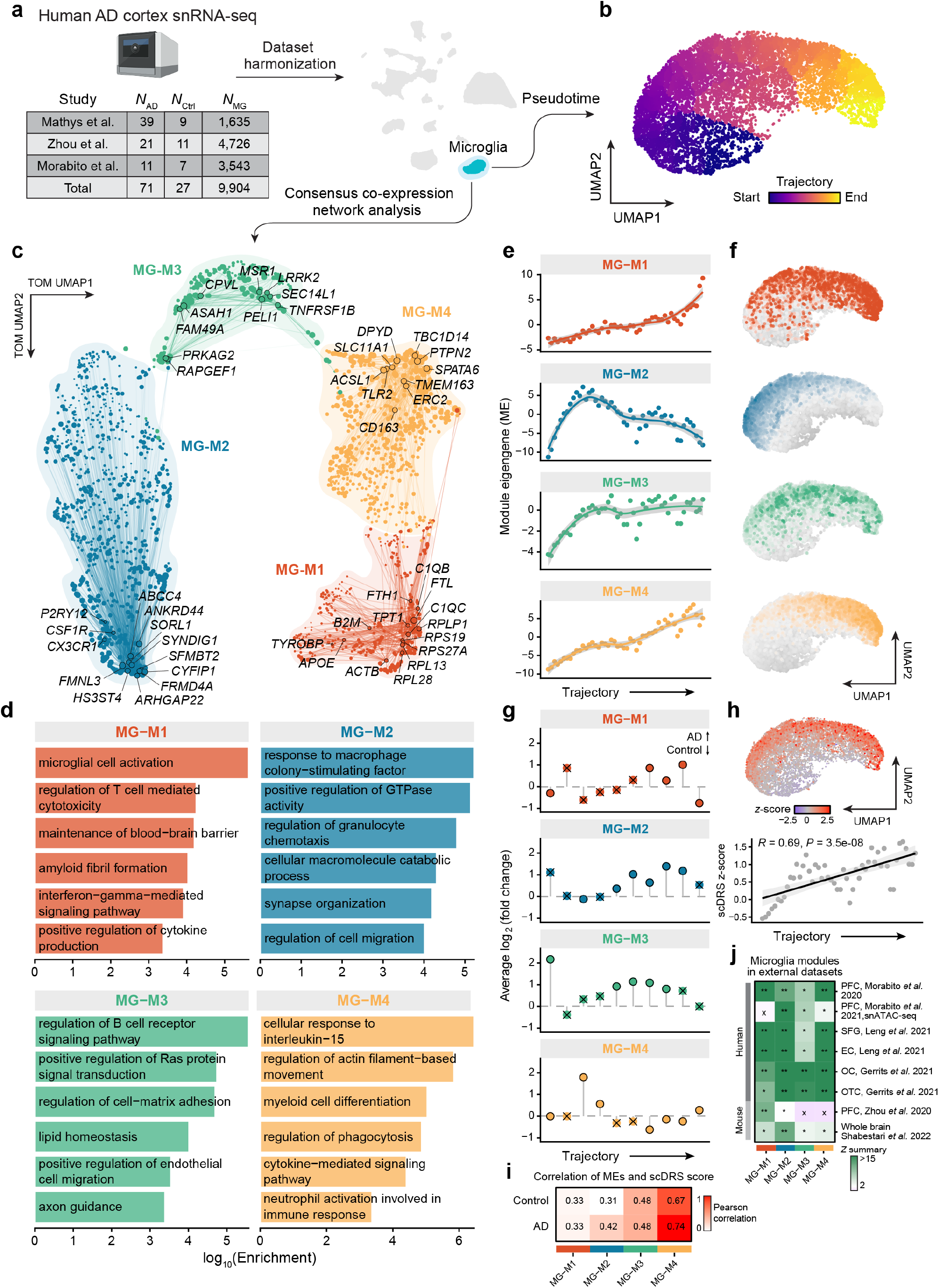
Consensus network analysis of microglia in Alzheimer’s disease. **a.** Left: table showing the number of samples and the number of microglia nuclei from published AD snRNA-seq datasets used for co-expression network analysis. Right: Integrated UMAP plot of nuclei from three snRNA-seq datasets. **b.** UMAP plot of microglia, colored by Monocle 3 (24) pseudotime assignment. **c.** UMAP plot of the microglia co-expression network. Each node represents a single gene, and edges represent co-expression links between genes and module hub genes. Point size is scaled by kME. Nodes are colored by co-expression module assignment. The top 10 hub genes per module are labeled, as well as additional genes of interest. Network edges were downsampled for visual clarity. **d.** Selected gene ontology (GO) terms enriched in co-expression modules. Bar plots show the log-scaled enrichment of each term. **e.** Module eigengenes (MEs) as a function of pseudotime, points are averaged MEs in 50 pseudotime bins of equal size. Line represents loess regression with a 95% confidence interval. **f.** Microglia UMAP colored by ME. **g.** Differential module eigengene (DME) results in ten pseudotime bins of equal size. For each pseudotime bin, we performed DME analysis between cells from AD (positive fold change) and control samples. × symbol indicates that the test did not reach significance (Wilcoxon rank sum test Bonferroni-adjusted *P*-value > 0.05). **h.** Top: Microglia UMAP colored by AD single-cell disease relevance score (scDRS) (48)*Z*-score. Bottom: scDRS *Z*-score as a function of pseudotime, points are averaged scDRS *Z*-scores in 50 pseudotime bins of equal size. Line represents linear regression with a 95% confidence interval. **i.** Heatmap of Pearson correlations of MEs and scDRS *Z*-scores, split by cells from AD and control samples. **j.** Heatmap showing module preservation Z-summary statistics for validation datasets. Abbreviations denote the following brain regions: Prefrontal cortex (PFC), superior frontal gyrus (SFG), entorhinal cortex (EC), occipital cortex (OC), occipitotemporal cortex (OTC). **: highly preserved (Z ≥ 10: highly preserved); *: moderately preserved (10 > *Z* ≥ 5); x: not preserved (*Z* < 5).

Classical markers of homeostatic microglia, such as *CSF1R, CX3CR1*, and *P2RY12* were members of MG-M2, while known DAM genes including *APOE, TYROBP*, and *B2M* were members of MG-M1. GO term enrichment analysis associated MG-M2 with homeostatic microglia functions such as cell migration, synapse organization, and response to colony-stimulating factor, contrasting disease-related processes enriched in MG-M1 including amyloid fibril formation, microglial activation, maintenance of blood-brain barrier, and cytokine production (Fig. 5d, Supplementary Table 18). Together, this suggests that MG-M1 comprises the gene network underlying disease-associated microglial activation in AD, while MG-M2 represents the network of homeostatic microglia genes. The MEs for MG-M1 and MG-M2 display opposing patterns throughout the microglia pseudotime trajectory, contextualizing this trajectory as the transcriptional shift from homeostatic microglia (start) to a DAM-like cell state (end) (Figs. 5e, f). Furthermore, DME analysis revealed significant changes in these modules between AD and control brains in evenly spaced windows throughout the microglia trajectory (Fig. 5g, Wilcoxon rank sum test Bonferroni-adjusted *P*-value < 0.05, Supplementary Table 19). Coexpression networks behave as functional biological units, therefore we reason that the hub genes and other members of MG-M1 represent candidates for an expanded set of human DAM genes including *ACTB, TPT1*, and *EEF1A1*.

Aside from modules MG-M1 and MG-M2 which contained well-known microglia gene signatures, we also identified modules MG-M3 and MG-M4 containing genes associated with key microglial processes like axon guidance, phagocytosis, and myeloid cell differentiation (Fig. 5c, d). *CD163*, a hub gene of MG-M4, is known to be involved in the breakdown of the blood-brain barrier (45, 46). The trajectory of MG-M4, containing *CD163* as a hub gene, was consistent with that of DAM-like module MG-M1, and was enriched for processes including phagocytosis, myeloid cell differentiation, and neutrophil activation (Fig. 5d), therefore it is possible that MG-M4 represents an alternative microglial activation module (47). We performed single-cell polygenic risk enrichment for AD risk in the microglia trajectory (39, 48), and identified a significant increase throughout the trajectory, revealing an enrichment of AD genetic risk SNPs in DAMs (Fig. 5h, Methods, Supplementary Table 20). We show that expression of the these modules were significantly correlated with AD genetic risk (Pearson correlation *P*-value < 0.05), with the strongest correlation in alternative activation module MG-M4 (Fig. 5i).

To ensure that these microglial modules were reproducibile across other datasets and in mouse models of AD, we performed module preservation analysis (20) (Methods, Fig. 5j). We projected the microglial consensus modules into a dataset of the PFC in aged human samples (49), the superior frontal gyrus (SFG) and entorhinal cortex (EC) in AD samples (50), the occipital cortex (OC) and the occipitotemporal cortex (OTC) in human AD samples (51), the PFC from 5xFAD mice (11), and whole brain samples from 5xFAD mice (23) (Fig. 5h). Additionally, we projected these modules into a snATAC-seq dataset of the PFC in human AD (12), using gene activity (52) as a proxy for gene expression from chromatin accessibility data. These module preservation tests showed the microglia consensus modules were broadly preserved and reproducible across brain regions and in mouse models of AD, providing further support that this network is relevant in AD biology and microglial activation.

### Projecting network modules from bulk RNA-seq cohorts into relevant single-cell datasets

hdWGCNA allows for interrogating co-expression modules inferred from a given reference dataset in a query dataset. Modules can be projected across datasets by computing module eigengenes in the query dataset, and preservation of the network structure can be assessed via statistical testing (20). For example, modules can be projected between different species to link transcriptomic changes between mouse models and human disease patients, or modules can be projected across data modalities from single-cell to spatial transcriptomics to provide regional context to cellular niches.

To date, it remains cost-prohibitive for most researchers to perform high dimensional -omics studies of large patient cohorts, but there are numerous large-scale disease-relevant bulk RNA-seq datasets containing thousands of samples from consortia such as ENCODE (53), GTEx (54), and TCGA (55). By projecting co-expression modules derived from bulk RNA-seq patient cohorts into single-cell datasets, we can layer disease-related information onto the single-cell dataset and attribute cell-state specific expression patterns to the bulk RNA-seq data. We demonstrate projecting modules in this manner using co-expression modules from two bulk RNA-seq studies of AD (13, 49) as the references and an AD snRNA-seq dataset (12) as the query. These studies both used AD samples and controls from the same patient cohorts (Religious Orders Study and Memory and Aging Project, Mayo clinic, Mount Sinai School of Medicine) (56–58), but they took unique approaches for co-expression network analysis. The AMP-AD study from Wan et al. (13) performed network analysis separately from each brain region, while in our previous study (49) we performed consensus network analysis across the different brain regions. We projected these modules into a snRNA-seq dataset of AD and control samples from the PFC (Fig. 6a), and we found distinct cell-type specific expression patterns based on their MEs (Figs. 6b, c). This analysis demonstrates hdWGCNA’s ability to transfer co-expression information across datasets to uncover otherwise unseen biological insights.

**Fig. 6.**
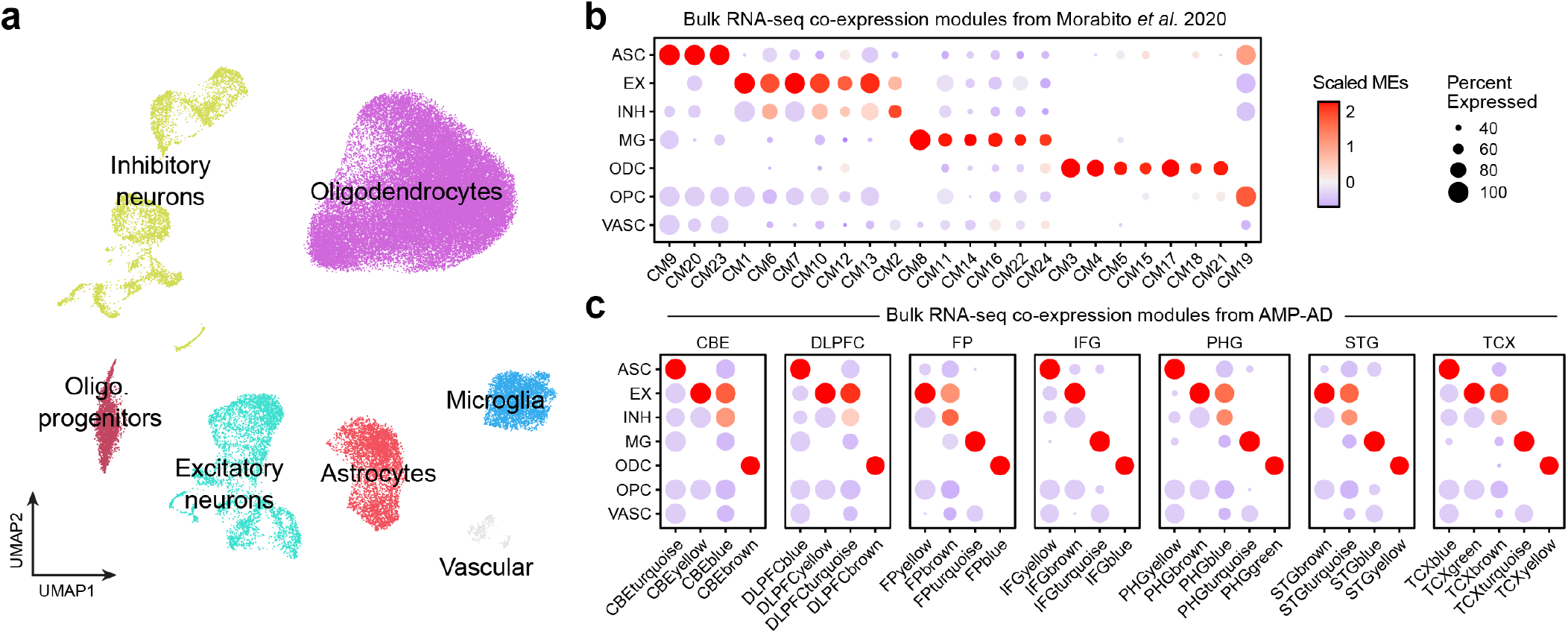
Projecting bulk RNA-seq co-expression modules into a single-cell dataset. **a.** UMAP plot of 57,950 nuclei from a snRNA-seq dataset of the human PFC from AD and control brains (12). Cells are colored by major cell type assignment. **b.** Multi-region consensus co-expression modules from Morabito *et al*. 2020 (49) bulk RNA-seq analysis projected into the snRNA-seq dataset as in panel a. **c.** Co-expression modules from the AMP-AD bulk RNA-seq dataset (13) projected into the snRNA-seq dataset as in panel a. CBE: cerebellum; DLPFC: dorsolateral prefrontal cortex; FP: frontal pole; IFG: inferior frontal gyrus; PHG: parahippocampal gyrus; STG: superior temporal gyrus; TCX: temporal cortex.

## Discussion

Classical bioinformatic approaches like differential gene expression analysis are useful for finding individual genes that are altered in a particular disease or condition of interest, but they do not provide information about the broader context of these genes in specific pathways or regulatory regimes. For example, biological processes like development or regeneration require coordination of distinct sets of genes in certain cell types with spatial specificity. Therefore to understand these complex processes, we must look beyond individual genes. hdWGCNA was developed to provide a succinct methodology for investigating systems-level changes in the transcriptome in single-cell or ST datasets. We designed hdWGCNA to be highly modular, allowing for multiscale analyses of different cellular or spatial hierarchies in a technology-agnostic manner.

In this study, we demonstrated that hdWGCNA is compatible with single-cell and ST datasets and can be easily adapted for novel transcriptomics approaches like ScISOrSeq. Co-expression networks have been successful for analyzing bulk proteomics datasets in human disease samples (59, 60), and we expect that hdWGCNA could be swiftly adapted for single-cell and spatial proteomics datasets as the technology matures and becomes more widely available (61). hdWGCNA includes built in functions to leverage external biological knowledge sources to provide insight for co-expression networks, for example by comparing gene modules to functional gene sets such as disease-associated genes from GWAS expression quantitative trait loci (eQTLs), or transcription factor target genes. Unlike other network analysis pipelines such as SCENIC (62) or CellChat (63), hdWGCNA is a purely unsupervised approach and does not require prior knowledge or databases in their inference procedure. The co-expression information computed by hdWGCNA can be easily retrieved from the Seurat object to facilitate custom downstream analyses beyond the hdWGCNA package. hdWGCNA allows for comparisons between experimental groups via differential module eigengene testing and module preservation analysis, which allowed us to identity inhibitory neuron modules that were dysregulated in ASD and enriched for ASD genetic risk genes, and microglial modules that were dysregulated in AD and enriched for DAM genes. Our network analyses of the ASD and AD datasets shows that hdWGCNA is capable of uncovering expanded disease-relevant gene sets via the interaction partners of known disease-associated genes like the ASD SFARI genes or the AD DAM genes. We demonstrated that the co-expression networks inferred by hdWGCNA were highly reproducible in unseen datasets, indicating that this is a robust methodology which reflects the underlying biology of the system of interest rather than picking up on technical artifacts. Further, hdWGCNA sheds new light on previously identified co-expression networks and gene modules by allowing modules to be projected from a reference dataset to a query dataset. The hdWGCNA R package directly extends the familiar Seurat pipeline and the SeuratObject data structure, enabling researchers to rapidly incorporate network analysis into their own workflows, going beyond cell clustering and differential gene expression analysis towards systems-level insights.

## ACKNOWLEDGEMENTS

Funding for this work was provided by UCI and UCI MIND start-up funds, National Institute on Aging grants 1RF1AG071683, U54 3U54AG054349-04S2, 3U19AG068054-02S, Adelson Medical Research Foundation funds, and an American Federation of Aging Research young investigator awarded to V.S. This work utilized the infrastructure for high-performance and high-throughput computing, research data storage and analysis, and scientific software tool integration built, operated and updated by the Research Cyberinfrastructure Center (RCIC) at the University of California, Irvine (UCI). We thank Anoushka Joglekar for providing the ScISOrSeq dataset.

## AUTHOR CONTRIBUTIONS

S.M. and V.S. conceptualized this study. The manuscript was written by S.M. F.R., and E.M. with assistance and approval from all authors. S.M. developed the hdWGCNA R package. S.M. and F.R. designed the structure of the hdWGCNA R package. S.M. collected, processed, and performed network analysis on publicly available sequencing datasets. F.R. performed bioinformatics analysis of the ScISOrSeq dataset. N.R. performed polygenic risk analysis of the integrated microglia snRNA-seq dataset.

## DATA AVAILABILITY

All data used in this study are publicly available, specified in Table 1.

## CODE AVAILABILITY

The hdWGCNA R package is available on GitHub at (https://github.com/smorabit/hdWGCNA), with documentation and tutorials available at (https://smorabit.github.io/hdWGCNA). All code used for data analysis in this paper is available on GitHub at (https://github.com/smorabit/hdWGCNA_paper).

## Methods

### Bootstrapped aggregation of single cell transcriptomes to form metacells

Single-cell gene expression datasets typically contain many more zero valued entries than non-zero valued entries, meaning that these datasets are sparse. We formally define the **sparsity** of a gene expression matrix in equation 1. Given an un-normalized counts matrix *X* with *N_g_* genes and *N_c_* cells, sparsity is the sum of all zero valued elements.

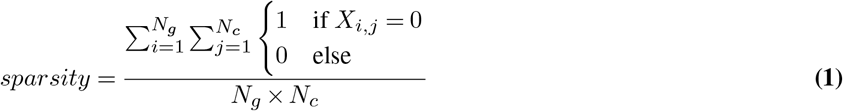

Complementing sparsity, the density of a single gene expression matrix is the sum of all non-zero valued elements, such that *density* = 1 – sparsity. A matrix is considered sparse if *sparsity* > 0.5. Conventional single-cell gene expression assays yield sparse gene expression matrices. In general, correlations of sparse vectors may lead to downstream conclusions that are not robust or reproducible. Thus, as part of the hdWGCNA workflow, we propose a bootstrapped aggregation (bagging) algorithm to construct a gene expression matrix *M* with considerably reduced sparsity prior to performing co-expression network analysis. Zero valued entries in a gene expression matrix have both biological and technical origins (64), and it is important to prioritize preserving relevant biological signals while reducing technical noise. For example, a biological zero may be attributed to a gene that is only expressed in a given cell population, whereas a technical zero may arise from low sequencing depth.

We define the set of unique cell barcodes *C* and the set of unique genes *G* such that ║*C*║ = *N_c_* and ║*G*║ = *N_g_*. Transcriptomically similar cells are identified in a dimensionally-reduced representation *D* of the gene expression matrix *X* using the *k*-nearest neighbors (KNN) algorithm (65), yielding *N_c_* sets of *k* cells. Inherently, there is overlap between these *N_c_* sets of *k* neighboring cells, and we include a parameter *m* to control for the maximum allowable overlap. Cells are uniformly randomly sampled from *C*, and gene expression signatures from *X* are aggregated (sum or average) with their *k* nearest neighbors. A cell is skipped if its neighbors have too much overlap with the set of neighbors from previously selected cells, in order to reduce redundancy in the downstream metacell expression matrix. The cell sampling loop converges when there are no more cells that satisfy the *m*, or when the number of target metacells *t* has been reached, yielding a metacell gene expression matrix *M*. Sparsity of the input and output matrices *X* and *M* are computed to check that sparsity is reduced throughout this process. This metacell bagging algorithm is implemented as part of the hdWGCNA R package in the `ConstructMetacells` function, and the pseudocode for this algorithm is defined in Algorithm 1. We denote a vector containing the elements of the *i*-th row of a matrix as *M_i*_* and a vector containing the elements of the *i*-th column as *M_*i_*.

#### Algorithm 1 ConstructMetacells

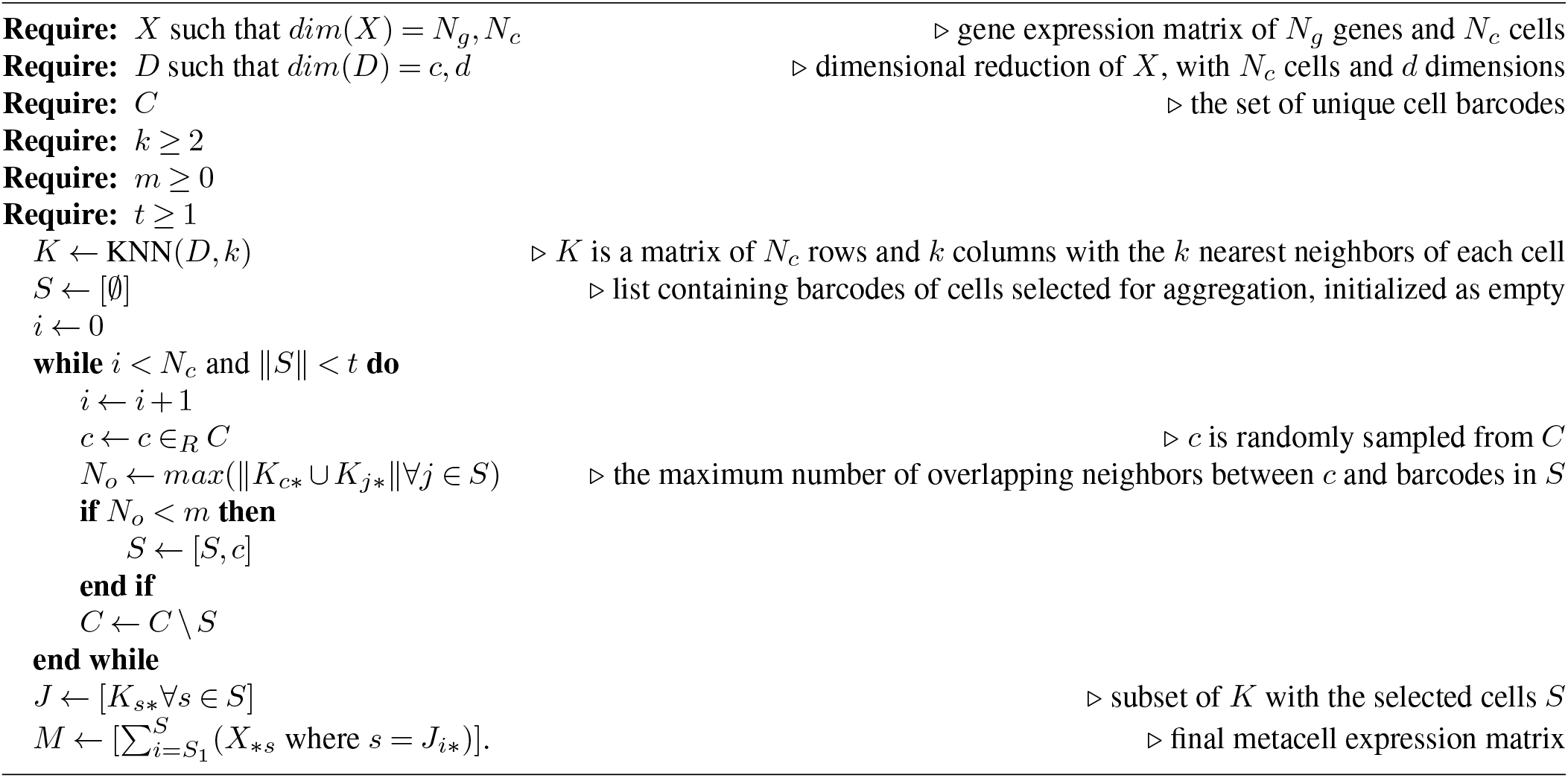

### Aggregation of neighboring spatial transcriptomic spots to form metaspots

Sequencing-based ST approaches such as the 10× Genomics Visium platform also yield sparse transcriptomic profiles, thus introducing the same potential pitfalls as single-cell data for co-expression network analysis. To alleviate these issues, we sought to develop a data aggregation approach similar to our metacell algorithm. This approach leverages spatial coordinates rather than the dimensionality-reduced representation (Fig. S5). For each ST spot, we obtain a list of physically neighboring spots. We then devise a grid of “principal spots”, which are evenly spaced spots throughout the input tissue which serve as anchor points for aggregating neighboring spots. Each principal spot and its neighbors are aggregated into one metaspot, with at most seven spots merging into one metaspot and at most two overlapping spots between metaspots. We implemented this procedure as part of the hdWGCNA R package in the `MetaspotsByGroups` function. Similar to the `MetacellsByGroups` function, the user may specify groups within the Seurat object to perform the aggregation, such that metacells would only be grouped within the same tissue slice, anatomical region, or other annotation. For all downstream analysis with hdWGCNA, the metaspot expression dataset can be used in place of the metacell expression matrix.

### Computing co-expression networks

Following metacell or metaspot construction, hdWGCNA constructs co-expression networks and identifies gene modules, building off of the WGCNA workflow (1, 66–69). The gene-gene adjacency matrix *A* is computed by taking the pairwise correlation of genes in *G* in the metacell expression matrix *M*, or in a subset of *M* for a specified cell population. Consider the gene expression vectors *x_i_* = *M_i*_* and *x_j_* = *M_j*_* for an arbitrary pair of genes (*i,j*) ∈ *G*, we compute the signed correlation as:

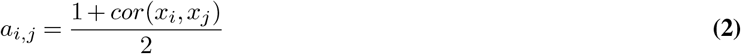

Note that *a_i,j_* is a linear transformation that retains the sign of the correlation while satisfying 0 ≤ *a_i, j_* ≤ 1. We define *A* as a symmetric adjacency matrix of size *N_g_ × N_g_* containing the signed correlations *a_i,j_* for all pairs (*i, j*) ∈ *G* as in equation 2. In order to emphasize strong correlations, we raise the elements of *A* to a power *β*, and we refer to this as soft power thresholding.

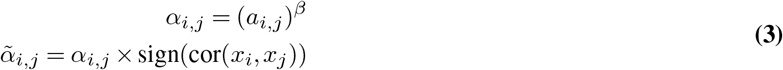

Now we have the gene-gene correlation raised to a power *β*, and an alternative metric 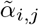 which also retains the sign of the correlation between these genes. The final co-expression network is then computed as a signed topological overlap matrix (TOM). The TOM describes shared neighbors between the a pair of genes (*i, j*). We define the signed TOM as

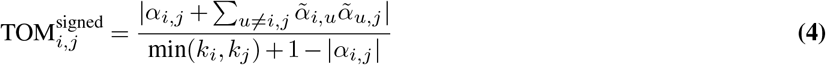

where *k_i_* and *k_j_* represent the connectivity between genes *i* and *j*

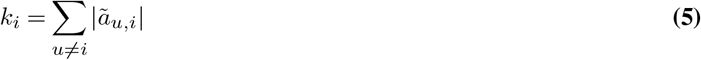

In the signed TOM, negative correlations serve to negatively reinforce the network connection, which is not the case in the unsigned TOM.

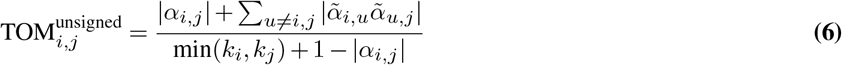

Genes are then grouped into modules based on the TOM network representation using the Dynamic Tree Cut algorithm (3), such that co-expression modules consist of genes with high topological overlap. Dynamic Tree Cut hierarchically clusters genes based on their dissimilarity in the TOM, denoted as DissTOM = 1 – TOM, thereby yielding a mapping between module assignments and gene names. The overall process transforming a metacell expression matrix *M* to a signed TOM co-expression network is implemented as part of the hdWGCNA R package in the `ConstructNetwork` function. Here we described the recommended workflow, using a signed adjacency matrix and a signed TOM, but `ConstructNetwork` can optionally construct unsigned or signed hybrid networks as well.

### Computing module eigengenes

Module eigengenes (MEs) are a convenient metric to summarize the gene expression of a given co-expression module. While the co-expression network was computed using the metacell expression matrix *M*, we compute MEs in the single-cell expression matrix *X*, thus yielding information about the activity of each module in each cell. The expression matrix for the *I*-th module consisting of genes *G*^(*I*^) ⊂ *G* is *X*^(*I*)^ = *X*_*G*(*I*),*_. The ME for module *I* is then computed by performing singular value decomposition (SVD), such that *X*^(*I*)^ = *UDV^T^*. Prior to running SVD, *X*^(*I*)^ must be scaled and centered, and we accomplish this using the Seurat function `ScaleData`. Importantly, `ScaleData` enables us to optionally perform regression to diminish the effects of selected technical covariates prior to computing MEs. The first column of *V*, containing the right-singular vectors 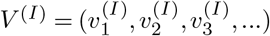 is the ME of module *I*.

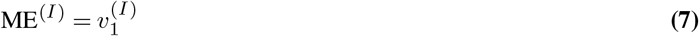

While SVD or other dimensionality reductions on a single-cell gene expression matrix contains critical biological information, technical artifacts are also present in these representations. There are many computational methods aiming to reduce technical effects in a reduced dimensional space, and these methods are often referred to as “batch-correction” or “integration” approaches (70). In particular, Harmony (71) is an algorithm well suited for correcting batch effects that may be present in a dimensionality-reduced single-cell expression dataset (70), and here we propose applying Harmony to MEs to maximize the biological information content of each ME. We implemented the ME computation algorithm, as defined in Algorithm 2, as part of the hdWGCNA R package in the function `ModuleEigengenes`.

#### Algorithm 2 ModuleEigengenes

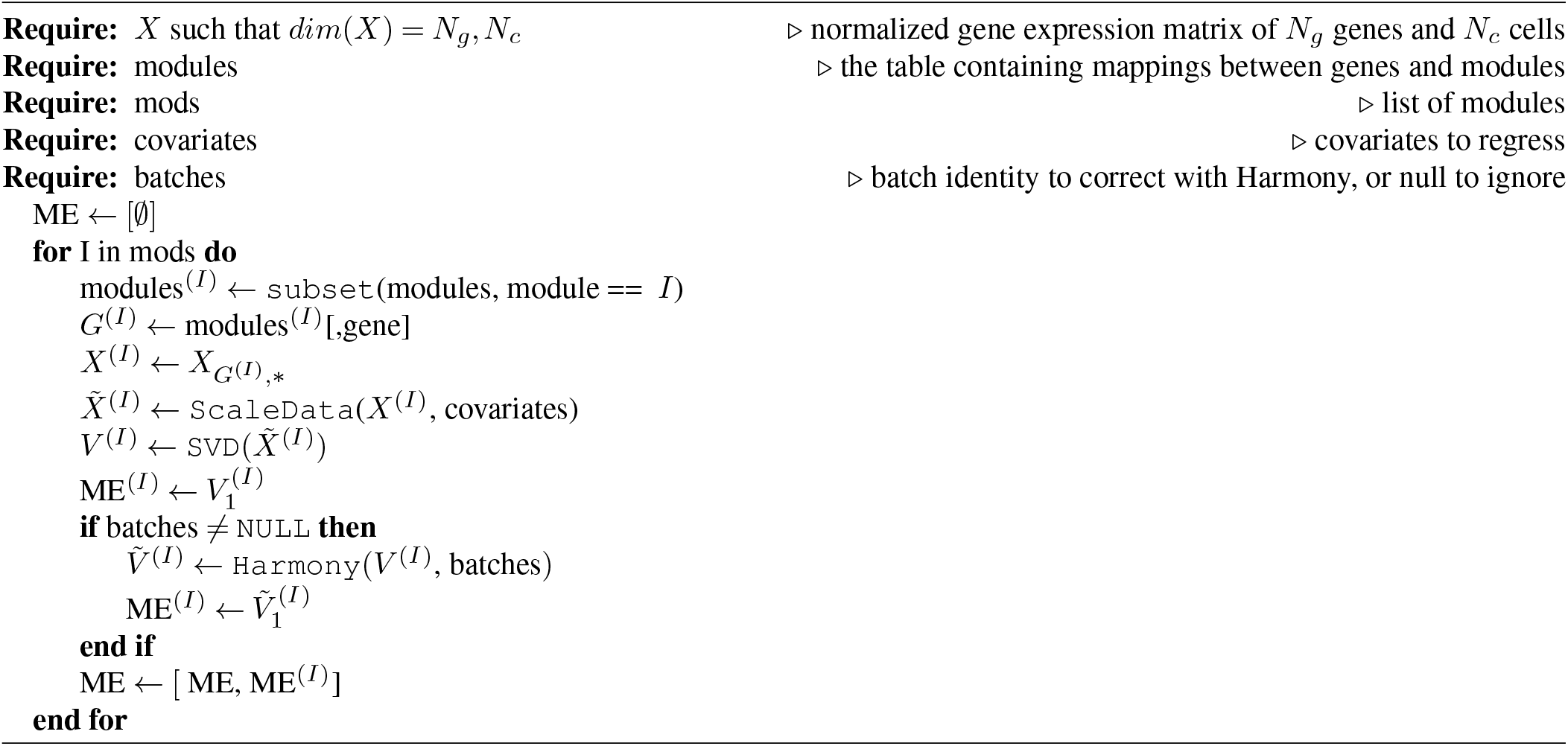

### Projecting co-expression modules in unseen data

In a typical hdWGCNA workflow, we perform metacell bagging, co-expression network analysis, module identification, and ME computation using the same single-cell gene expression dataset, starting from the expression matrix *X*. Given the module-gene assignment table derived from a reference dataset *X*, we can run the `ModuleEigengenes` algorithm on a query dataset *Y* where the genes in *Y* must be contained in the set of genes in *X* such that *G_Y_* ⊆ *G_X_*. We implemented this process in the hdWGCNA R package as the `ProjectModules` function. Importantly, we designed `ProjectModules` to be agnostic towards the data modality or species used in the reference and query datasets, thereby allowing for a host of comparative analyses. `ProjectModules` can facilitate cross-species analysis leveraging a table that maps gene symbols between two genomes. Modules can be projected into epigenomic data modalities such as single-cell assay for transposase accessible chromatin with sequencing (scATAC-seq) provided a measure of gene expression estimated from chromatin accessibility, such as Signac (52) gene activity or ArchR (72) gene scores. This approach can also be used to project modules from bulk expression datasets into single-cell or spatial transcriptomics datasets.

### Reprocessing published datasets

Table 1 details the different datasets used throughout this manuscript. We used several published datasets generated by our own group (12, 23, 49), and sequencing data was not re-downloaded for these studies. For all human snRNA-seq datasets, we applied a uniform processing pipeline to process each dataset starting from the raw sequencing data and resulting in an anndata object (73) containing UMI counts, normalized gene expression, cluster identities, and cell type annotations. Parameters used throughout this processing pipeline vary slightly between different datasets, and all parameters are noted in the data processing scripts in our github repository. For each biological replicate, we used the `kb count` function from kallisto | bustools (74) to psuedoalign raw sequencing reads to the reference transcriptome and quantify gene expression attributed to each cell barcode. The human reference transcriptome (GRch38) was obtained from the 10× Genomics website (version 2020-A, July 2020), and was re-formatted for use with kallisto | bustools using the `kb ref` function.

**Table 1.** Sequencing datasets used throughout this manuscript. Data was either obtained from the Sequence Read Archive (SRA), Synapse, a custom website (Parse Biosciences), the SeuratData R package, or directly from the authors, as denoted in the Data source column. Tissue abbreviations are the following; PFC: prefrontal cortex; EC: entorhinal cortex; SFG: superior frontal gyrus; HC: hippocampus; PBMCs: peripheral blood mononuclear cells.

| Study / Dataset | Data source | Species | Tissue | Technology | $N_{\text{replicates}}$ | $N_{\text{barcodes}}$ |
| --- | --- | --- | --- | --- | --- | --- |
| Mathys *et al.* 2019 (10) | syn18485175 | human | PFC | 10× V2 snRNA-seq | 48 | 48,905 |
| Velmeshev *et al.* 2019 (9) | PRJNA434002 | human | PFC | 10× V2 snRNA-seq | 54 | 121,451 |
| Morabito *et al.* 2020 (49) | authors | human | PFC | 10× V2 snRNA-seq | 5 | 35,771 |
| Nagy *et al.* 2020 (78) | GSE144136 | human | PFC | 10× V2 snRNA-seq | 34 | 83,849 |
| Zhou *et al.* 2020 (11) | syn21670836 | human | PFC | 10× V2 snRNA-seq | 32 | 82,366 |
| Zhou *et al.* 2020 (11) | syn21670836 | mouse | PFC | 10× V2 snRNA-seq | 6 | 38,034 |
| Gerrits *et al.* 2021 (51) | GSE148822 | human | OC | 10× V2 snRNA-seq | 18 | 203,932 |
| Gerrits *et al.* 2021 (51) | GSE148822 | human | OTC | 10× V2 snRNA-seq | 18 | 220,176 |
| Joglekar *et al.* 2021 (8) | authors | mouse | HC | ScISOrSeq | 1 | 6,832 |
| Leng *et al.* 2021 (50) | GSE147528 | human | EC | 10× V2 snRNA-seq | 10 | 44,465 |
| Leng *et al.* 2021 (50) | GSE147528 | human | SFG | 10× V2 snRNA-seq | 10 | 57,439 |
| Morabito *et al.* 2021 (12) | authors | human | PFC | 10× V3 snRNA-seq | 18 | 57,856 |
| Morabito *et al.* 2021 (12) | authors | human | PFC | 10× snATAC-seq | 20 | 132,623 |
| Shabestari *et al.* 2022 (23) | authors | mouse | brain | ParseBio snRNA-seq | 48 | 51,327 |
| 10× Genomics brain ST | SeuratData | mouse | brain | 10× Genomics Visium | 2 | 6,049 |
| ParseBio PBMCs | custom | human | PBMCs | ParseBio scRNA-seq | 24 | 965,363 |

For each of the UMI counts matrices, we used the `remove-background` function from cellbender (75) to simultaneously identify which barcodes corresponded to cells and to remove counts attributed to ambient RNA. We then used scrublet (76) to compute “doublet scores”, the likelihood of each barcode mapping to more than one cell. Counts matrices from each biological replicate in a given dataset are then merged into a single anndata object, and any relevant sample level meta-data (age, sex, disease status) was stored in the `adata.obs` table. We performed a percentile filtering of cells that were outliers from each dataset based on the number of UMI per cell the percentage of UMI attributed to mitochondrial genes per cell, and the doublet score. Filtering based on these criteria was performed in each sample, as well as dataset-wide. After filtering, downstream data processing steps were carried out with SCANPY (73). The UMI counts matrix was normalized with *ln*(CPM) using the functions `sc.pp.normalize_total` and `sc.pp.log1p`. Highly variable genes were identified using the function `sc.pp.highly_variable_genes`, and these genes are used as the features for downstream analysis steps such as principal component analysis (PCA). The normalized expression matrix was then scaled to unit variance and centered at zero using the function `sc.pp.scale`. PCA was performed on the scaled expression matrix using the function `sc.tl.pca`. Harmony (71) was used to correct the PCA matrix for batch effects using the function `sc.external.pp.harmony_integrate`. The harmonized PCA matrix was then used to construct a cell neighborhood graph using the function `sc.pp.neighbors`. The cell neighborhood graph was then used to compute a two-dimensional representation of the data with uniform manifold approximation and projection (19) using the function `sc.tl.umap`, and to group cells into clusters with Leiden clustering (77) using the function `sc.tl.leiden`. We inspected the gene expression signatures in each Leiden cluster for a panel of canonical cell-type marker genes in order to assign a cell-type label to each cluster, and to identify additional doublet clusters that may have escaped the previous filtering steps. The distribution of quality control metrics was inspected in each cluster. We filtered out cells belonging to clusters that displayed conflicting expression of cell-type marker genes, or were outliers in their quality control metrics. After filtering these low-quality clusters, we ran UMAP and Leiden clustering again, resulting in the final processed dataset.

### Iterative network analysis of major cell types in the human cortex

We performed an iterative co-expression network analysis of the major cell types (ASC, EX, INH, MG, ODC, OPC) in the human PFC snRNA-seq dataset from Zhou *et al*. (11), only including samples from control brains (36,671 cells and 36,601 genes). We retained genes that were expressed in at least 5% of cells for downstream analysis. Metacells were computed separately for each major cell type and each sample using the hdWGCNA function `MetacellsByGroups`, aggregating 25 cells per metacell. Further, we ran `MetacellsByGroups` while varying the *K* parameter in order to asses the resulting metacell expression matrix sparsity. For each cell type, we applied the following hdWGCNA commands with default arguments to perform network analysis: `TestSoftPowers`, `ConstructNetwork`, `ModuleEigengenes`, `ModuleConnectivity`, and `RunModuleUMAP`. We performed module preservation analysis (20) of the ODC co-expression modules in an external snRNA-seq dataset of the human PFC (12). Modules were projected from the reference to query dataset using the hdWGCNA function `ProjectModules`, and the module preservation test was performed using `ModulePreservation` with 100 permutations.

### Comparison with alternative metacell approaches

For the purpose of co-expression network analysis, we compared our metacell aggregation approach (Algorithm 1) with two alternative approaches, namely Metacell2 (15) and SEACells (17). We ran the three metacell approaches using the recommended settings on the same dataset, and then ran hdWGCNA on each of the resulting metacell expression matrices. We used a scRNA-seq of 6,800 CD34+ hematopoietic stem and progenitor stem cells included with the SEACells package, and we used the cluster annotations from the original study. Notably, SEACells and Metacell2 do not account for cell labels in their aggregation procedures, which may result in a number of metacells containing transcriptomes from differently labeled cells. For the hdWGCNA metacell algorithm, we aggregated 50 cells per metacell. For the three metacell expression matrices derived from the different algorithms, we performed co-expression network analysis with the standard hdWGCNA pipeline by sequentially running the following functions with default parameters: `TestSoftPowers`, `ConstructNetwork`, `ModuleEigengenes`, `ModuleConnectivity`, and `RunModuleUMAP`. With the same cluster settings, Dynamic Tree Cut recovered a different number of co-expression modules for the three methods (hdWGCNA: 16 modules; MC2: 13 modules; SEACells: 20 modules). We performed pairwise comparisons between the gene modules detected with each metacell approach using Fisher’s exact test to test module overlaps. Additionally, we performed rank-rank hypergeometric overlap (79) (RRHO) tests using the `RRHO` function from the R package RRHO (version 1.13.0) to compare the kME ranking between modules across methods. To compare MEs and Seurat module scores, we ran the `AddModuleScore` function, and computed Pearson correlations between each ME and each module score.

### Application of hdWGCNA to a one million cell scRNA-seq dataset

We obtained a publicly available scRNA-seq dataset from Parse Biosciences of 1M peripheral blood mononuclear cells (PBMCs) from 12 healthy donors and 12 Type-1 diabetic donors generated using the Evercode Whole Transcriptome Mega protocol. This analysis was performed on a compute cluster with 200 GB of memory and eight CPU cores. The UMI counts matrix and sample meta data was downloaded from Parse Biosciences’ website. We processed the counts matrix using SCANPY using a similar pipeline as described in the *Reprocessing published dataset* section. For quality control, we excluded cells with greater than 25% mitochondrial reads, greater than 5,000 genes, and greater than 25,000 counts. After dimensionality reduction with PCA, Harmony (71) batch correction, and Leiden clustering (77) (resolution=1), we annotated cell populations using PBMC marker genes obtained from Azimuth (7). We excluded clusters with conflicting cell-type markers as potential doublet populations, retaining a total of 965,363 cells and 26,862 genes for downstream analysis. The major cell compartments recovered in this analysis were similar to those reported by Parse Biosciences in their analysis, including as T-cells, B-cells, monocytes, dendritic cells, basophils, and plasmablasts. Following the SCANPY data processing, we wrote the individual components (counts matrix, cell meta-data, gene meta-data, dimensionality reductions, etc.) to disk so they could be loaded into R and assembled into a Seurat object.

We performed co-expression network analysis iteratively for the plasmablast, T-cell, B-cell, monocyte, and dendritic cell compartments using an hdWGCNA pipeline for each group (Fig. S4). Metacells were constructed separately for each sample and each cell cluster with the hdWGCNA function `MetacellsByGroups`, aggregating 50 cells per metacell. The metacell aggregation step had a runtime of 85 minutes and 59 seconds. For each cell population, we first subset the Seurat object for the cell population of interest and then performed the standard hdWGCNA pipeline by sequentially running the following functions with default parameters: `TestSoftPowers`, `ConstructNetwork`, `ModuleEigengenes`, `ModuleConnectivity`, and `RunModuleUMAP`. We note that for the largest cell population (T-cells, 555,417 cells), the runtime for the network construction step was 186 seconds.

### Spatial co-expression network analysis in the mouse brain

We collected the publicly available 10× Genomics Visium mouse brain dataset using the SeuratData R package. This dataset consists of an anterior and a posterior slice from a sagittal brain section, which we merged into a single Seurat object comprising 6,049 ST spots and 31,053 genes. We processed this dataset using the standard Seurat pipeline by sequentially running the following commands: `NormalizeData`, `FindVariableFeatures`, `ScaleData`, `RunPCA`, `FindNeighbors`, `FindClusters`, and `RunUMAP`. The top thirty PCs were used for Louvain clustering (80) and UMAP. While ST spots were clustered based on transcriptomic information alone, we were able to annotate them based on anatomical features.

Neighboring ST spots were aggregated into metaspots in the anterior and posterior slices using the hdWGCNA function `MetaspotsByGroups`. We retained genes expressed in 5% of spots for downstream analysis, totaling 12,355 genes. We tested for the optimal soft-power threshold *β* based on the the fit to a scale-free topology using the hd-WGCNA function `TestSoftPowers`. The co-expression network was constructed using all ST spots spanning both the anterior and posterior slices using the hdWGCNA function `ConstructNetwork` with the following parameters: networkType=“signed”, TOMType=“signed”, soft_power=5, deepSplit=4, detectCutHeight=0.995, minModuleSize=50, merge-CutHeight=0.2. Module eigengenes and eigengene-based connectivities were computed using the `ModuleEigengenes` and `ModuleConnectivity` functions respectively. This approach identified 12 spatial co-expression modules, and we visualized the spatial distributions of these modules by plotting their MEs directly onto the biological coordinates for each spot. The co-expression network was projected into two dimensions using UMAP with the hdWGCNA function `RunModuleUMAP`, and we used the top five hub genes (ranked by kMEs) as the input features for UMAP. We used the R package enrichR (81) (version 3.0) to perform enrichment analysis on the top 100 genes in each module ranked by kME using the following databases: GO_Biological_Process_2021, GO_Cellular_Component_2021, GO_Molecular_Function_2021, WikiPathway_2021_Mouse, and KEGG_2021_Mouse. We assessed the overlap between genes from these spatial co-expression modules and differentially expressed genes in each cluster from a recent snRNA-seq study of the whole mouse brain using Fisher’s exact test implemented in the R package GeneOverlap (version 1.26.0). Finally, we performed a separate network analysis on a subset of the ST dataset only containing the cortical layers 2-6, and we followed an identical hdWGCNA analysis pipeline to the full ST dataset for the cortical analysis.

### Isoform co-expression network analysis in the mouse hippocampus

We performed isoform co-expression network analysis in radial glia lineage cells (radial glia, astrocytes, ependymal cells, and neural intermediate progenitor cells) from mouse hippocampus ScISOrSeq dataset from Joglekar *et al*. (8)using the hdWGCNA R package. The gene-level counts matrix for this dataset was obtained from GEO (GSE15845), and the isoform-level counts matrix was obtained directly from the authors of the original study. We formatted this dataset as a Seurat object with an isoform-level expression assay and a gene-level expression assay. The standard Seurat processing pipeline was used on the gene-level expression assay, where we sequentially ran the functions `NormalizeData`, `FindVariableFeatures`, `ScaleData`, and `RunPCA` with default parameters. The dataset was projected into two dimensions by running UMAP on the PCA matrix with 30 components using the `RunUMAP` function. For all downstream purposes, the cell-type annotations from the original study were used.

Radial glia cells were selected for network analysis, and isoforms expressed in fewer than 1% of these cells were excluded, yielding a set of 2,190 cells and 10,375 isoforms from 4,770 genes. We constructed metacells separately for each cell type on the isoform-level expression assay using the hdWGCNA function `MetacellsByGroups` with *k* = 30. We performed a parameter sweep for the soft-power threshold *β* using the function `TestSoftPowers`. The isoform co-expression network was constructed using the `ConstructNetwork` function with the following parameters: networkType=“signed”, TOMType=“signed”, soft_power=5, deepSplit=4, detectCutHeight=0.995, minModuleSize=50, mergeCutHeight=0.5. This approach identified 11 isoform co-expression modules. Isoform-level module eigenisoforms were computed using the `ModuleEigengenes` function, and eigenisoform-based connectivity was computed using the `ModuleConnectivity` function with default parameters. We computed a semi-supervised UMAP projection of the co-expression network using the hdWGCNA function `RunModuleUMAP`, with the module labels and the top six hub isoforms (by kMEiso) per module as the input features. We used the enrichR to identify enriched pathways in each module ranked by using the following databases: GO_Biological_Process_2021, GO_Cellular_Component_2021, GO_Molecular_Function_2021, WikiPathway_2021_Mouse, and KEGG_2021_Mouse.

To assess isoform co-expression network dynamics throughout the cellular trajectories within the radial glia lineage, we performed pseudotime analysis using Monocle 3 (33) (version 1.0.0). We computed a UMAP of just radial glia lineage cells using the Monocle 3 function `run_umap`. A trajectory graph was built on this UMAP representation using the function `learn_graph`, and pseudotime was calculated with the function `order_cells` using the radial glia cells as the starting point. We split the pseudotime trajectory into three lineages based on the distinct cell fates (astrocyte, neuronal, and ependymal). We grouped cells into 50 evenly-sized bins throughout each trajectory, and we applied loess regression to the average module eigenisoform of each module in these bins to inspect the dynamics of each module throughout development. We wrote a custom script to generate a GTF of isoform models output from the ScISOrSeq pipeline. To visualize expressed isoforms, we plotted isoforms from this GTF on the UCSC genome browser as well as in Swan (25).

### Co-expression analysis of inhibitory neurons in autism spectrum disorder (ASD)

We selected inhibitory neurons from the Velmeshev *et al*. (9) human ASD snRNA-seq dataset for co-expression network analysis. Of the 121,451 cells in this dataset, 20,249 were labeled as inhibitory neurons based on marker gene expression profiles. We retained 11,194 genes which were expressed in at least 10% of cells from any cluster, and had non-zero variance in the inhibitory neuron population. Metacell transcriptomic profiles were constructed separately for each of the 54 samples and each cell type using the hdWGCNA function `MetacellsByGroups`, aggregating 50 cells into one metacell. We selected a soft-power threshold *β* = 9 based on the parameter sweep performed with the `TestSoftPowers` function. The co-expression network was computed with the `ConstructNetwork` function with the following parameters: networkType=“signed”, TOMType=“signed”, soft_power=9, deepSplit=4, detectCutHeight=0.995, minModuleSize=50, mergeCutHeight=0.2. Module eigengenes were computed using the `ModuleEigengenes` function, and we applied Harmony (71) to correct MEs based on sequencing batch. Eigengene-based connectivity for each gene was computed using `ModuleConnectivity`. The co-expression network was embedded in two dimensions using UMAP with the `RunModuleUMAP` function with the top five genes (ranked by kMEs) per module as the input features. Distributions of MEs were compared between ASD and control samples for each inhibitory neuron subpopulation using a two-sided Wilcoxon rank sum test with the R function `wilcox.test`. We used the enrichR (81) to perform enrichment analysis on the top 100 genes in each module ranked by kME using the following databases: GO_Biological_Process_2021, GO_Cellular_Component_2021, GO_Molecular_Function_2021, WikiPathway_2021_Human, and KEGG_2021_Human. Furthermore, we computed the overlap between co-expression modules and ASD-associated genes from the SFARI Gene database using the R package GeneOverlap, which calculates the overlap between sets of genes using Fisher’s exact test.

### Consensus co-expression network analysis of microglia in Alzheimer’s disease (AD)

We performed consensus co-expression network analysis of microglia in Alzheimer’s disease using three published snRNA-seq datasets (10–12). The individually processed datasets were merged into a single Seurat object comprising 189,127 nuclei, and the datasets were integrated into a common dimensionally-reduced space using PCA and Harmony (71). We retained all nuclei labeled microglia for network analysis based on expression of canonical marker genes such as *CSF1R* (9,904 nuclei), and genes expressed in at least 5% of microglia from any of the three studies were retained (7,900 genes). Metacells were constructed in groups of cells based on AD diagnosis status and study of origin, aggregating 25 cells per metacell. Within hdWGCNA, we used the `SetMultiExpr` function to create a list of expression matrices containing the selected genes and metacells for the three studies. We performed a separate parameter sweep for the three expression matrices using the hdWGCNA function `TestSoftPowerConsensus`, ensuring that we used an appropriate *β* value for each dataset (Mathys *et al*.: *β* = 6, Zhou *et al*.: *β* = 8, Morabito & Miyoshi et al: *β* = 6). The consensus co-expression network was contructed using the hdWGCNA function `ConstructNetwork` using the `consensus=TRUE` option. Individual TOMs were computed for each dataset, and they were scaled based on the 80th percentile in order to alleviate different statistical properties specific to each dataset rather than the underlying biology. A consensus TOM was computed by taking the element-wise minimum of the individual TOMs from each dataset. Therefore, large topological overlap values between two genes, which indicate a strong co-expression relationship, are supported across all three datasets in the consensus TOM. We performed hierarchical clustering on the consensus TOM, and we used the Dynamic Tree Cut algorithm (3) was used to identify consensus co-expression modules based on the hierarchy. Module eigengenes were computed using the `ModuleEigengenes` function, and we applied Harmony (71) to correct MEs based on the dataset of origin. Eigengene-based connectivity for each gene was computed using `ModuleConnectivity`. We visualized the network using UMAP with the top ten hub genes (ranked by kMEs) per module as the input features, annotating the hub genes and known disease-associated microglia genes (43). We used the enrichR (81) to perform enrichment analysis on the top 100 genes in each module ranked by kME using the following databases: GO_Biological_Process_2021, GO_Cellular_Component_2021, GO_Molecular_Function_2021, WikiPathway_2021_Human, and KEGG_2021_Human.

We sought to model the transcriptional dynamics governing the shift between homeostatic and activated microglia in AD, therefore we performed pseudotime analysis using Monocle 3 (33) to build a continuous trajectory of microglia cell states. A trajectory graph was built on the microglia UMAP using the function `learn_graph`, and pseudotime was calculated with the function `order_cells`. We oriented the start of pseudotime based on the expression of homeostatic microglia marker genes, such as *P2RY12, CX3CR1*, and *CSF1R*. We grouped cells into 50 evenly-sized bins throughout each trajectory, and we applied loess regression to the average module eigengene of each module in these bins to inspect the dynamics of each module throughout the microglia trajectory.

To link the integrated microglia snRNA-seq dataset with polygenic risk of disease for individual cells, we used the scDRS python package (version 1.0.0) (48). This pipeline takes 1) a set of putative disease genes derived from GWAS summary statistics and 2) a scRNA-seq dataset as inputs, and outputs disease enrichment statistics for a given disease (raw and normalized disease scores, cell-level scDRS p-value, and *Z*-scores converted from the p-values). GWAS summary statistics of 74 diseases and complex traits supplied by scDRS were utilized as gene sets, among which a gene set by Jansen *et al*. (39) provided the set of genes associated with AD. We then visualized the AD scDRS *Z*-scores in the integrated AD microglia trajectory, and we correlated the scDRS score with the trajectory using a Pearson correlation.

We performed module preservation (20) analysis in a variety of external datasets from human and mouse (11, 12, 23, 49–51) to test for the reproducibility of the consensus AD microglia modules in the microglia population from each dataset. We used the hdWGCNA function `ProjectModules` to compute module eigengenes for the consensus AD microglia modules for each query dataset. The module preservation test was performed using the hdWGCNA function `ModulePreservation` with 100 permutations, and we reported the preservation *Z*-summary statistics in a heatmap. For the Morabito & Miyoshi *et al*. snATAC-seq dataset, we used the gene activity (52) representation as a gene-level summary of chromatin accessibility in order to assess the module preservation at the epigenomic level.

### Analysis of bulk RNA-seq co-expression modules in single-cell data

We projected gene co-expression modules from two bulk RNA-seq studies of AD (13, 49) into a published snRNA-seq study of AD to assess their expression patterns within various cell populations. While both of these studies used the samples from the same bulk RNA-seq cohort, the set of modules from Morabito *et al*. 2020 (49) was based on a consensus network analysis across six brain regions while the other set of modules from the AMP-AD study (13) were constructed separately for seven different brain regions. Module eigengenes were computed for each of these bulk RNA-seq modules in the snRNA-seq dataset using the hdWGCNA function `ProjectModules`, using Harmony to correct MEs based on sequencing batch. We visualized the MEs of the projected modules in the snRNA-seq dataset using the Seurat function `DotPlot`.

**Fig. S1.**
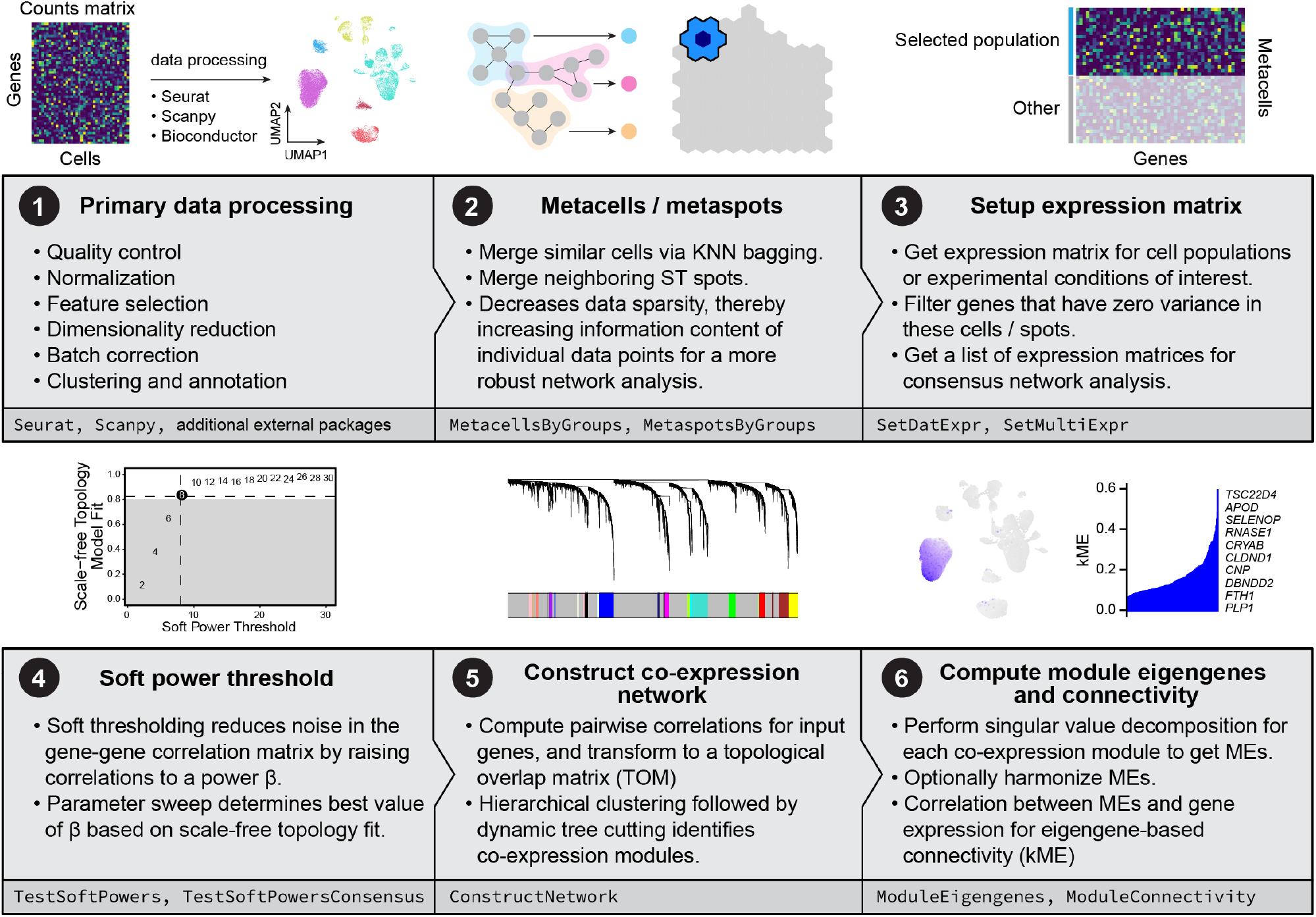
Schematic of the hdWGCNA workflow. **1.** Prior to analysis with the hdWGCNA R package, the input single-cell or spatial dataset must be fully processed. This includes quality control, data normalization, feature selection, dimensionality reduction, batch correction (if needed), and clustering. These steps can be done using popular packages such as Seurat (5–7) or SCANPY (73). Regardless of the pipeline used, the dataset must be formatted as a Seurat object prior to running hdWGCNA. **2.** The functions `MetacellsByGroups` and `MetaspotsByGroups` are used to aggregate transcriptomically similar cells into metacells and spatially proximal spots into metaspots respectively. **3.** hdWGCNA requires the user to explicitly specify the expression matrix that will be used for network analysis using the function `SetDatExpr`. For consensus network analysis, a list of expression matrices for each dataset or condition is specified with `SetMultiExpr`. **4.** Different values for the soft-power threshold *β* are tested using the functions `TestSoftPowers` and `TestSoftPowersConsensus`. The gene-gene correlation adjacency matrix is computed and raised to a power *β* as a soft threshold, and the degree distribution of this augmented is fit to a power law distribution to assess the scale-free topology. **5.** Co-expression network computation and gene module detection is performed in one step using the function `ConstructNetwork`. **6.** Module eigengenes (MEs) are computed using the `ModuleEigengene` function, optionally allowing for regression and harmonization of covariates such as sequencing batch. Eigengene-based connectivity (kME) is computed for each gene using the `ModuleConnectivity` function.

**Fig. S2.**
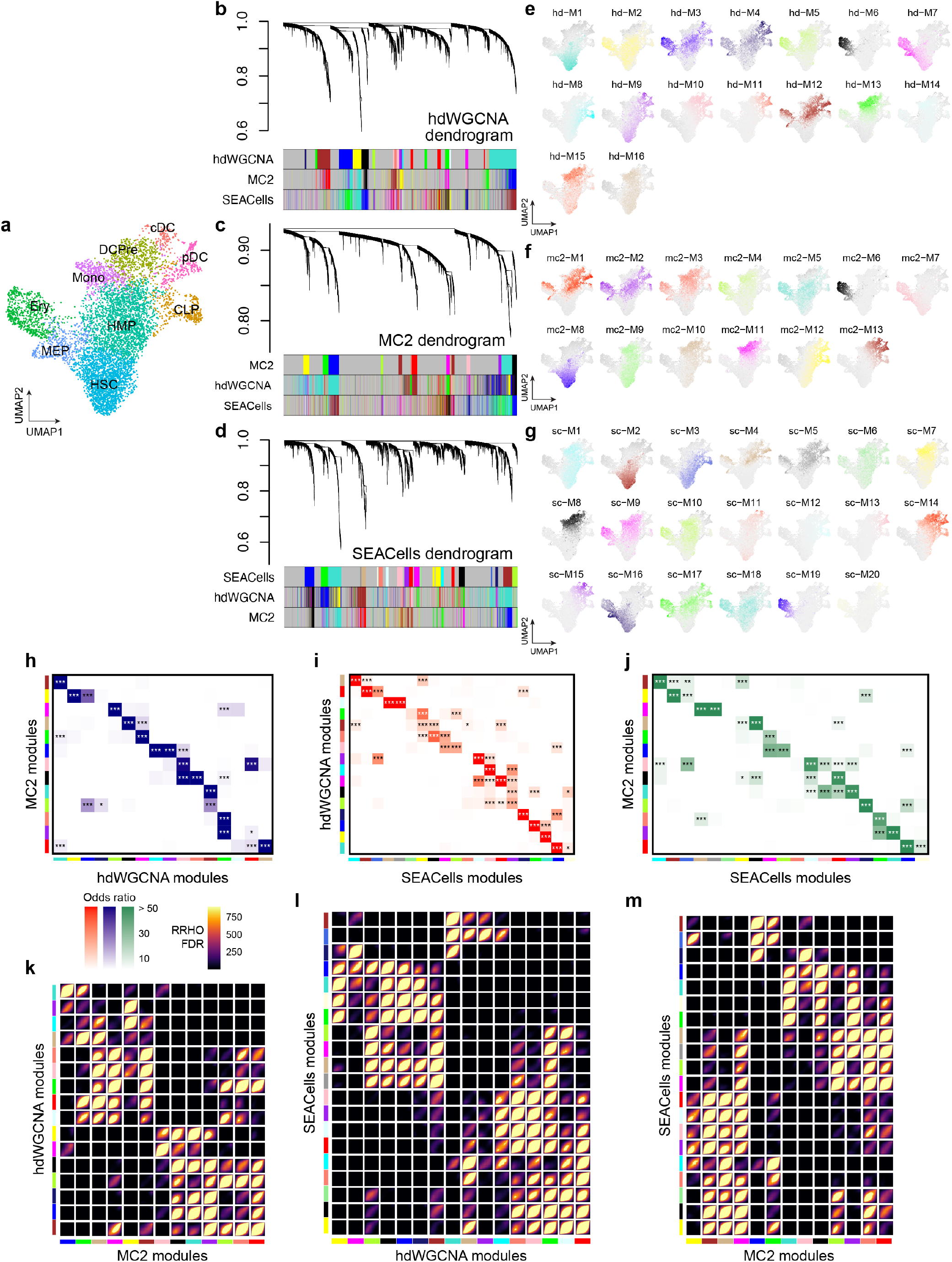
Comparison of metacell algorithms for co-expression network analysis. **a.** UMAP plot of the 6,800 CD34+ hematopoietic stem and progenitor stem cells scRNA-seq dataset (17) colored by annotations from the original study. **b-d.** hdWGCNA dendrograms for the co-expression networks constructed with the hdWGCNA (**b**), MC2 (**c**), and SEACells (**d**) algorithms. Module assignments are shown below the dendrograms **e-g.** UMAP plots as in (**a**) colored by MEs for the co-expression modules derived from the different metacell approaches. **h-j.** Module overlap comparisons between the different methods. Test was performed using Fisher’s exact test, and we report the odds ratio and FDR corrected p-values. **k-m.** Rank-rank hypergeometric overlap (RRHO) (79) heatmaps comparing the ranks of kMEs for pairs of modules derived from the expression matrices from the different metacell algorithms.

**Fig. S3.**
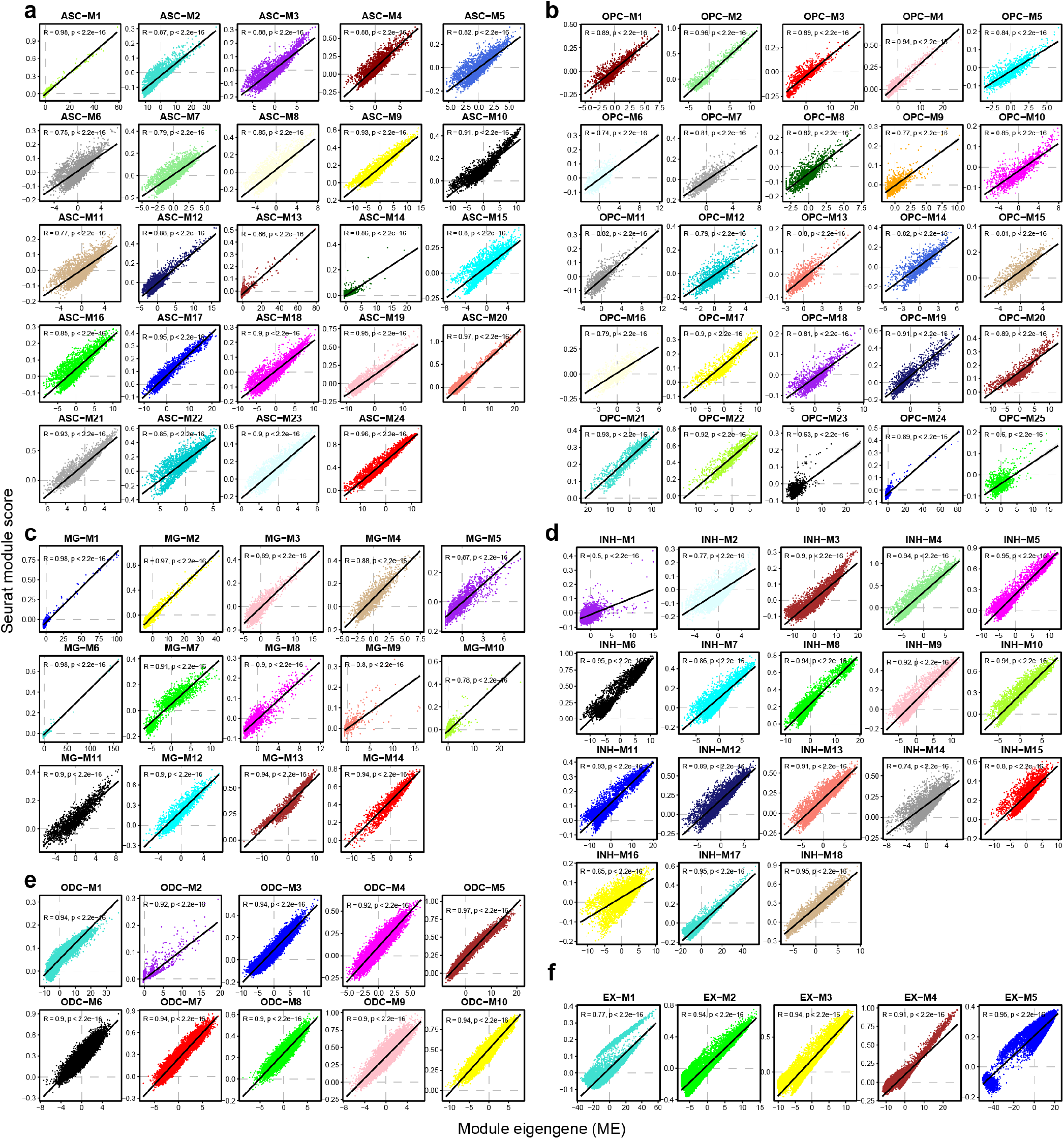
Correlation of module eigengenes and Seurat module scores. For each module in the human PFC snRNA-seq dataset (11), we computed Seurat module scores using the *`AddModuleScore`* function, and correlated with module eigengenes (MEs). We visualized the results as scatter plots with linear regression lines (95% confidence interval shown in grey) for ASC (**a**), OPC (**b**), MG (**c**), INH (**d**), ODC (**e**), and EX (**f**).

**Fig. S4.**
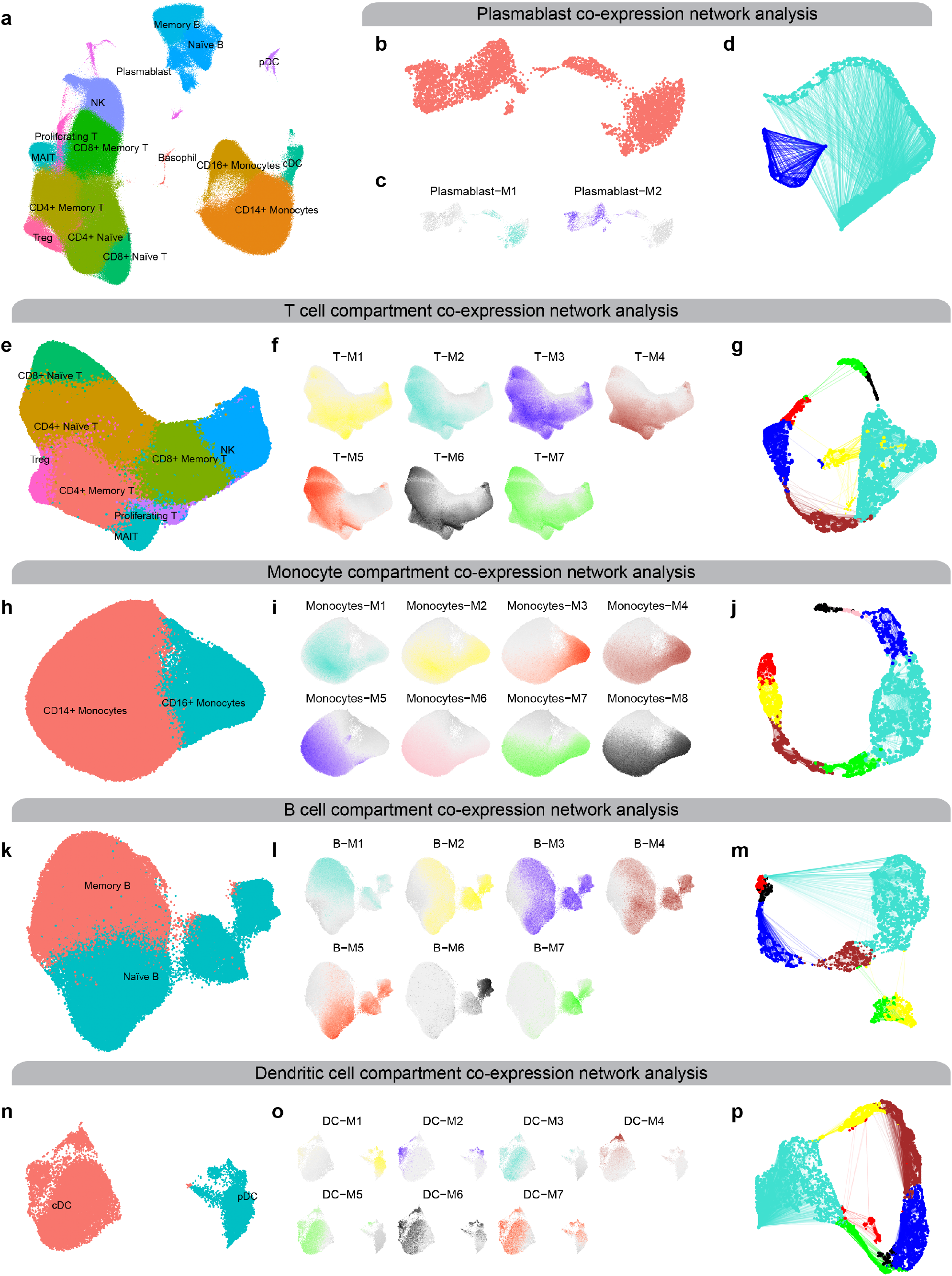
Iterative co-expression network analysis of major cell compartments in the Parse Biosciences 1M PBMC dataset. **a.** UMAP plot of 965,363 PBMCs from 12 healthy donors and 12 Type-1 diabetic donors profiled with the Parse Biosciences Evercode Whole Transcriptome Mega protocol. Cells are colored by cell-type annotations. **b,e,h,k,n.** Individual UMAP plots computed for each cell compartment separately for each cell compartment. **c,f,i,l,o.** UMAP plots for each cell compartment colored by module eigengenes (MEs). **d,g,j,m,p** UMAP plots of the co-expression networks for each cell compartment, colored by gene module assignment. Nodes represent genes and edges represent co-expression links. Network edges were downsampled for visual clarity.

**Fig. S5.**
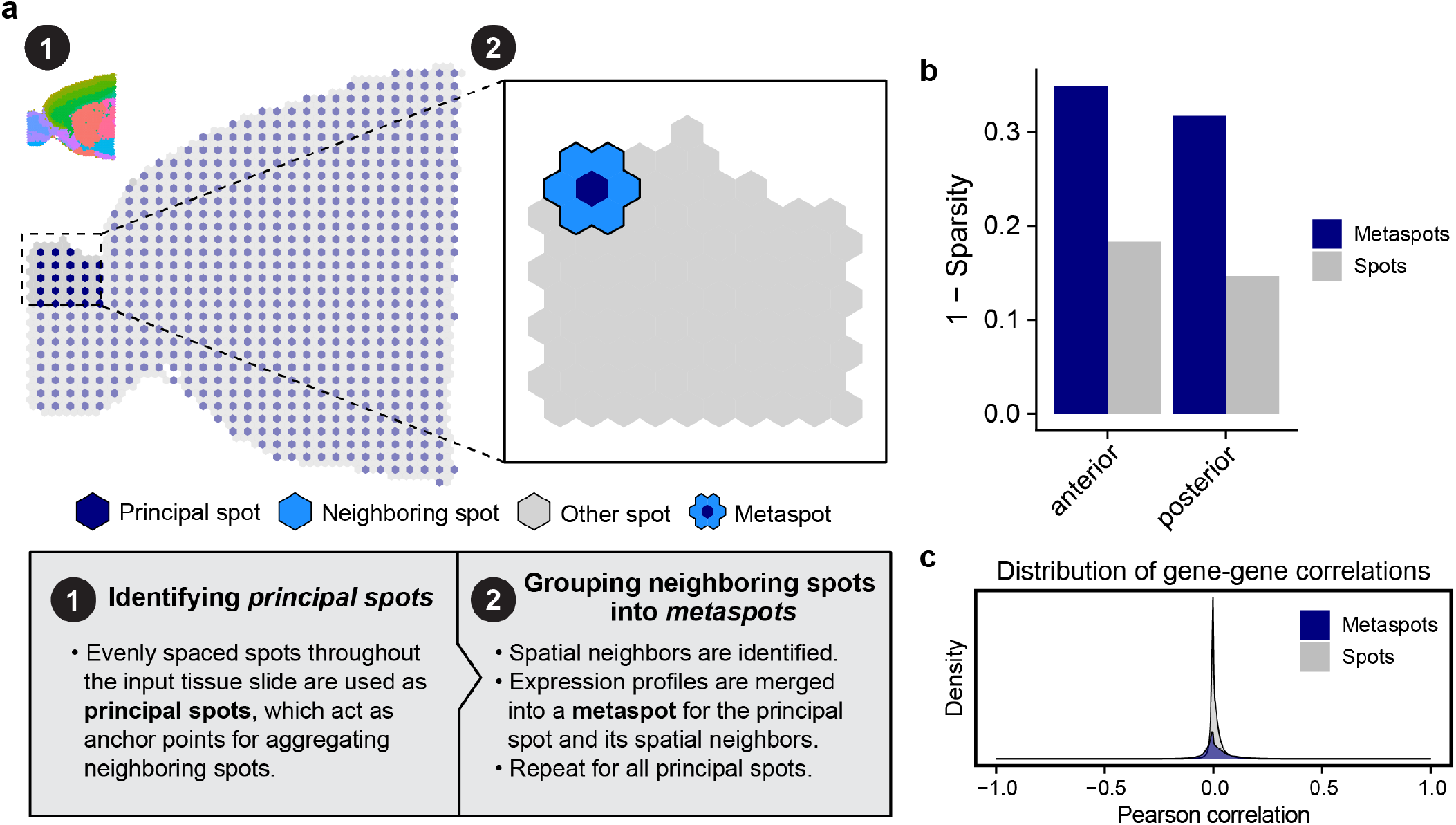
Metaspot aggregation for co-expression network analysis in spatial transcriptomics. **a.** Schematic representation of the metaspot construction process. A grid of evenly spaced *principal spots* are specified throughout the given input ST section. The expression values for each principal spot and its direct neighbors are merged into a single metaspot expression profile. This procedure yields a metaspot expression matrix for the given input ST section. **b.** Expression matrix density (1 - sparsity) for the ST and metaspot expression matrices in the anterior and posterior mouse brain samples. **c.** Density plot showing the distribution of pairwise Pearson correlations between genes from the ST expression matrix and the metaspot expression matrix.

**Fig. S6.**
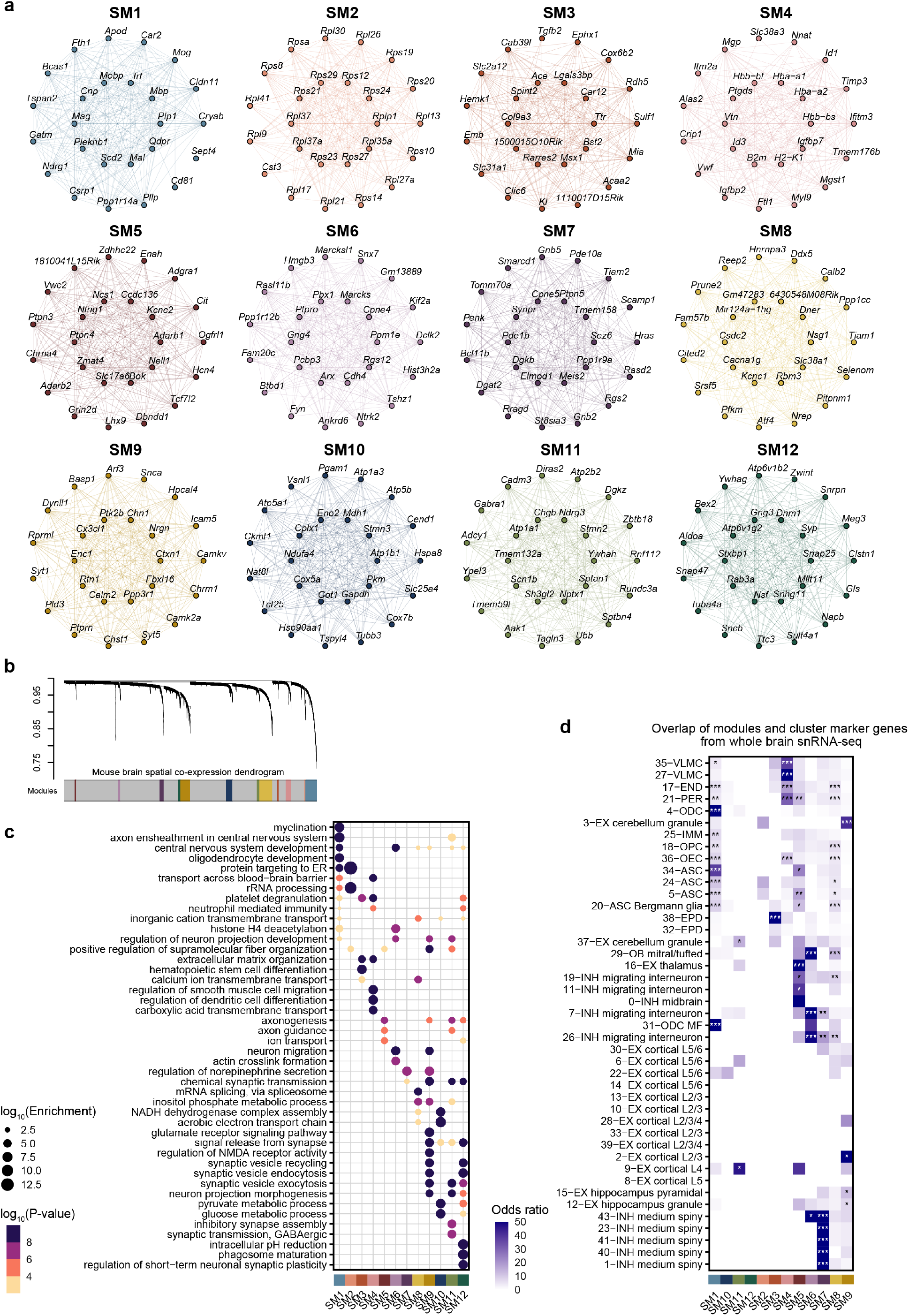
Mouse brain spatial transcriptomics co-expression network analysis. **a.** Hub gene networks for each spatial co-expression module. The top 25 hub genes ranked by kME are visualized. Nodes represent genes, and edges represent co-expression links. **b.** Dendrogram showing the hierarchical clustering of genes into co-expression modules based on the topological overlap matrix (TOM). **c.** Dot plot showing selected GO term enrichment results for each co-expression module. **d.** Heatmap showing gene overlap tests (Fisher’s exact test) comparing the spatial co-expression modules to differentially expressed genes (DEGs) in each cluster from a whole mouse brain snRNA-seq dataset (23).

**Fig. S7.**
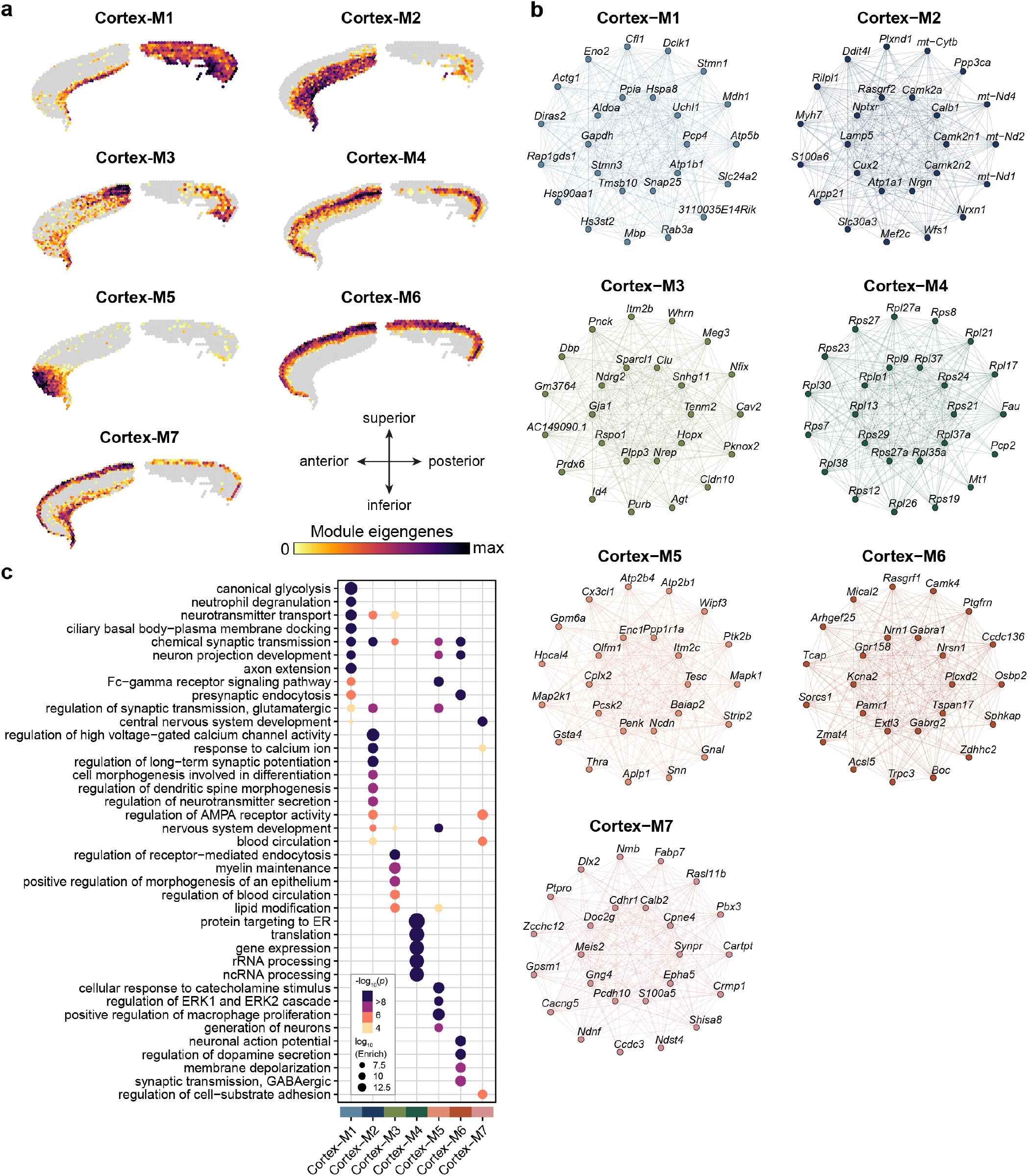
Mouse brain spatial transcriptomics co-expression network in cortical layers 2-6. **a.** ST samples colored by module eigengenes (MEs) for the seven cortical spatial co-expression modules. Grey color indicates a ME values less than zero. **b.** Hub gene networks for each cortical co-expression module. The top 25 hub genes ranked by kME are visualized. Nodes represent genes, and edges represent co-expression links. **c.** Dot plot showing selected GO term enrichment results for each co-expression module.

**Fig. S8.**
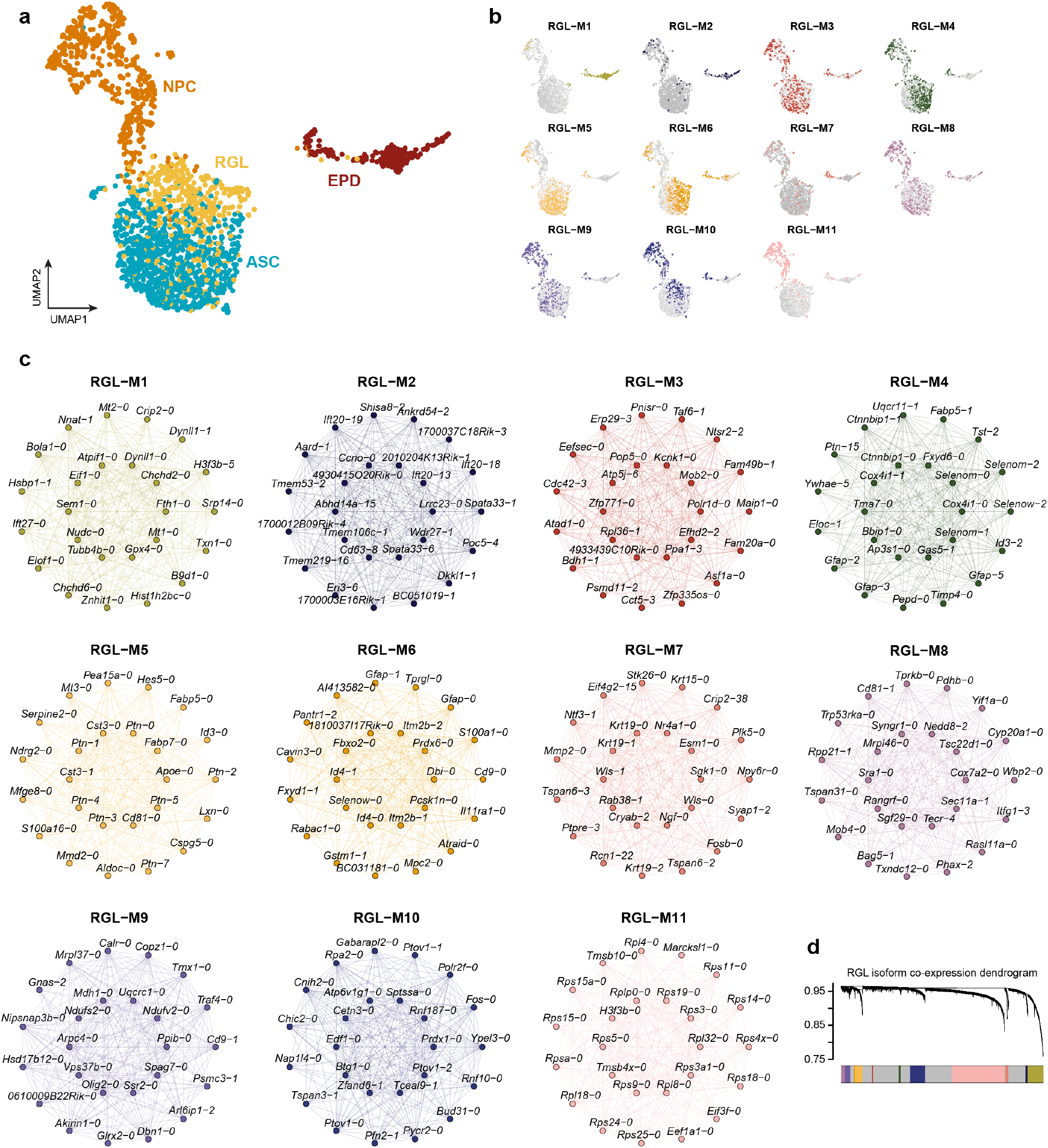
Isoform co-expression network analysis in the mouse hippocampus. **a.** UMAP plot of the radial glia lineage clusters from the mouse hippocampus ScISOrSeq dataset (8). **b.** UMAP plots as in **a.** colored by MEiso for the eleven isoform co-expression modules. **c.** Hub isoform networks for each radial glia lineage co-expression modules. The top 25 hub genes ranked by kMEiso are visualized. Nodes represent isoforms, and edges represent co-expression links. **d.** Dendrogram showing the hierarchical clustering of isoforms into co-expression modules based on the topological overlap matrix (TOM).

**Fig. S9.**
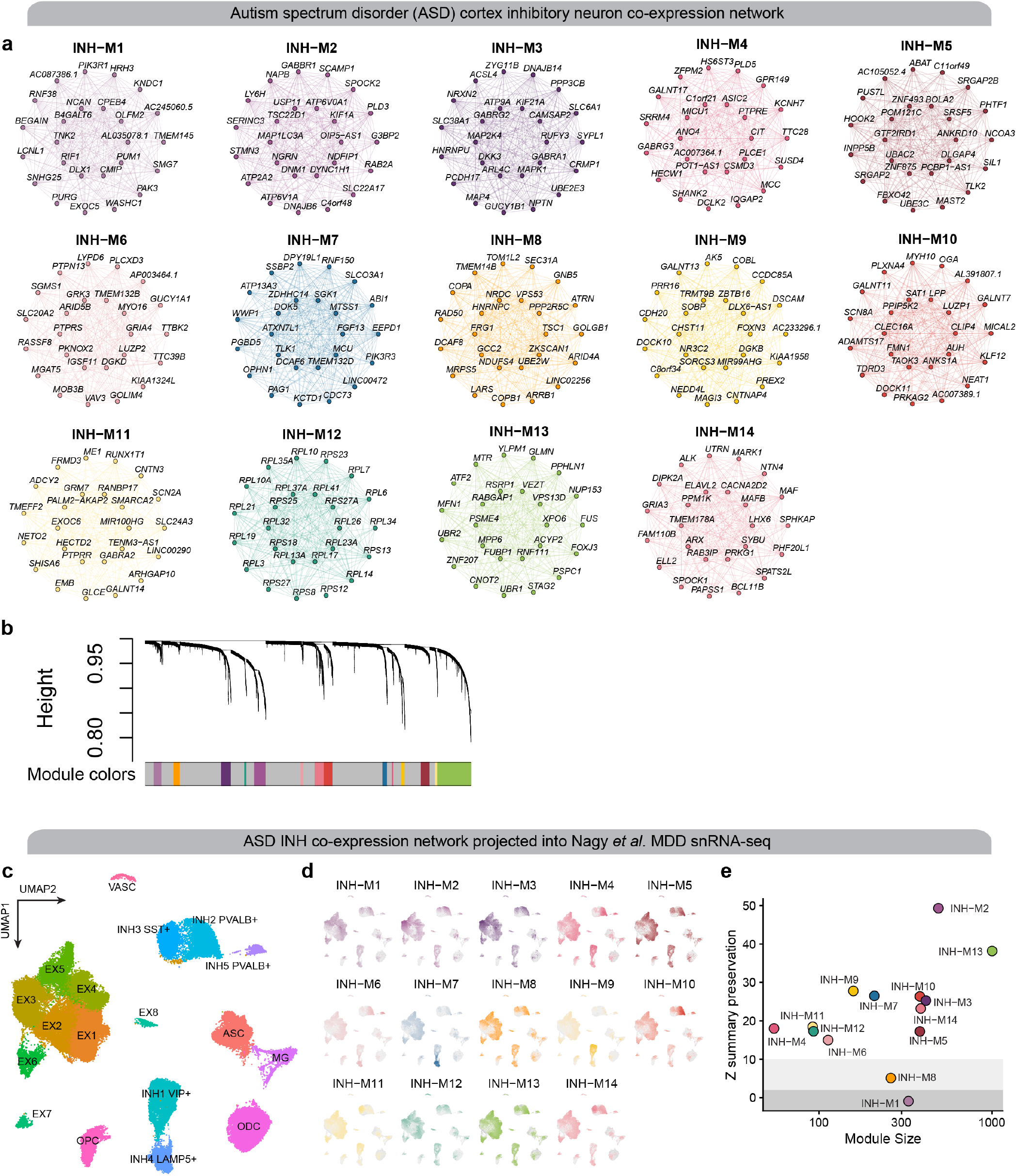
Co-expression network analysis of inhibitory neurons in Autism spectrum disorder. **a.** Hub gene networks for each inhibitory neuron co-expression modules. The top 25 hub genes ranked by kME are visualized. Nodes represent genes, and edges represent co-expression links. **b.** Dendrogram showing the hierarchical clustering of genes into co-expression modules based on the topological overlap matrix (TOM). **c.** UMAP plot of the snRNA-seq dataset of human major depressive disorder (MDD) (78). Cells are colored by cluster annotations. **d.** UMAP plots of the MDD dataset as in **c.** colored by the MEs projected from the ASD dataset. **e.** Module preservation statistics for the ASD inhibitory neuron modules in the inhibitory neuron population from the MDD dataset. *Z*-summary preservation < 2 indicates no evidence of module preservation, *Z*-summary preservation < 10 indicates moderate evidence of module preservation, and *Z*-summary preservation > 10 indicates high evidence of module preservation

**Fig. S10.**
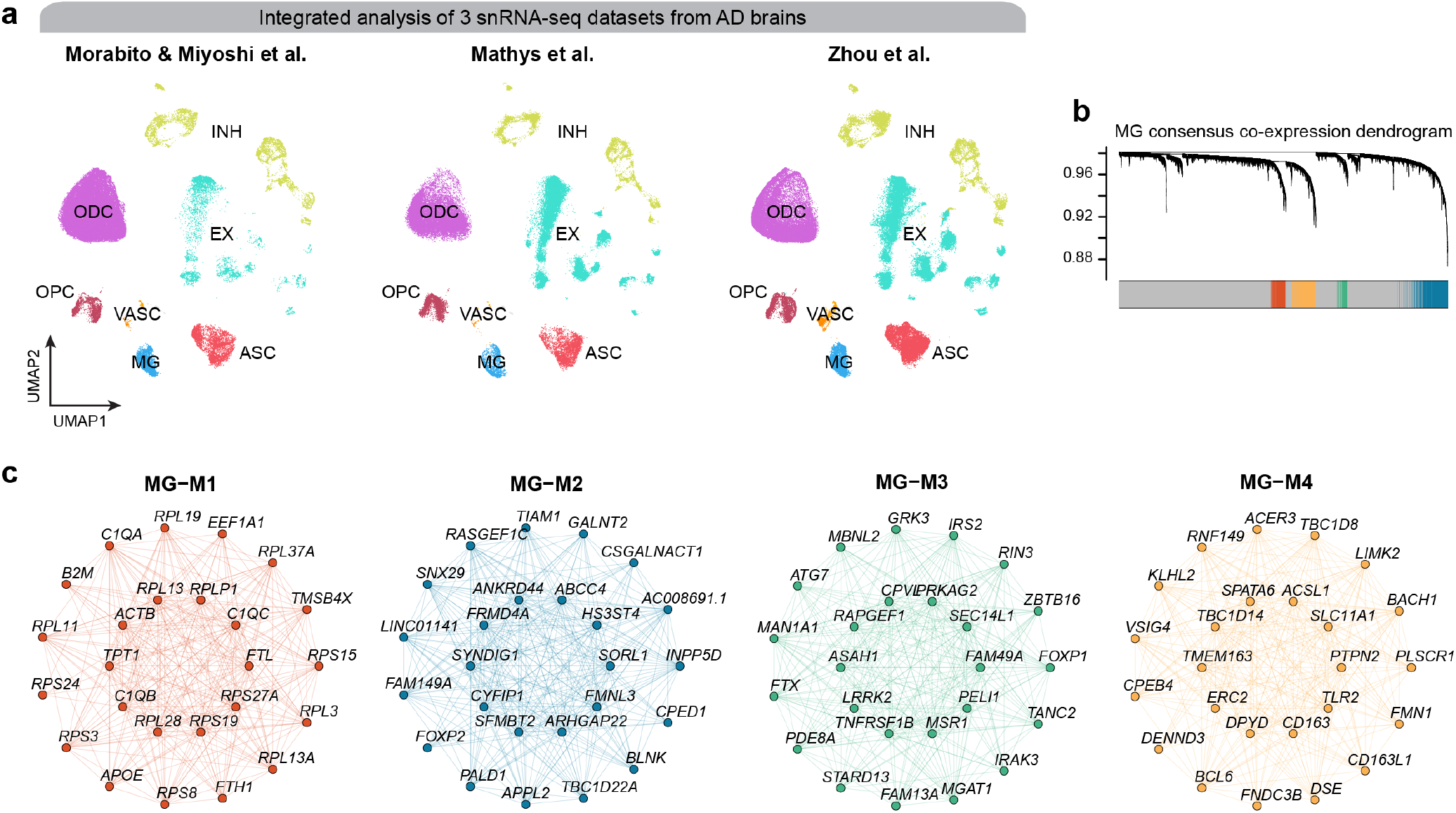
Consensus co-expression network analysis of microglia in Alzheimer’s disease. **a.** UMAP plot of the integrated dataset from three AD snRNA-seq studies (10–12), colored by cell type assignment. **b.** Dendrogram showing the hierarchical clustering of genes into co-expression modules based on the consensus topological overlap matrix (TOM) from the three snRNA-seq datasets. **c.** Hub gene networks for each microglia consensus co-expression modules. The top 25 hub genes ranked by kME are visualized. Nodes represent genes, and edges represent co-expression links.

